# TAK1 operates at the primary cilium in non-canonical TGFB/BMP signaling to control heart development

**DOI:** 10.1101/2024.05.06.592628

**Authors:** Canan Doganli, Daniel A. Baird, Yeasmeen Ali, Oskar Kaaber Thomsen, Enrique Audain, Line Jessen, Pauline Munck Truelsen, Johanne Bay Mogensen, Maria Schrøder Holm, Kateřina Apolínová, Lorenzo Buttò, Maria Diamanti, Jindřiška Leischner Fialová, Emma M. Wade, Stephen P. Robertson, Lotte Bang Pedersen, Laurent Argiro, Fabienne Lescroart, Marc-Phillip Hitz, Søren Tvorup Christensen, Lars Allan Larsen

**Affiliations:** Department of Cellular and Molecular Medicine, University of Copenhagen, Copenhagen, Denmark; Department of Biology, University of Copenhagen, Copenhagen, Denmark; Department of Medical Genetics, Carl von Ossietzky University, Oldenburg, Germany; ZeClinics SL, Badalona, Spain; Faculty of Health and Life Sciences, Universitat Pompeu Fabra, Barcelona, Spain; Department of Women’s and Children’s Health, Dunedin School of Medicine, University of Otago, New Zealand; Aix Marseille Université, INSERM, MMG U1251, Marseille, France

**Keywords:** Heart development, cardiomyogenesis, congenital heart disease, primary cilia, TAK1, MAP3K7, TAB2, PKA-Cα, PRKACA

## Abstract

Transforming Growth Factor-Beta-Activated Kinase 1 (TAK1/MAP3K7), along with its upstream regulators TAK1-Binding Protein 2 (TAB2) and the catalytic alpha-subunit of Protein Kinase A (PKA-Cα/PRKACA), has been identified as a pivotal player in regulation of developmental processes. Haploinsufficiency of TAB2 causes Congenital Heart Disease (CHD) and rare variants in PKA-Cα and TAK1 cause cardioacrofacial dysplasia (CAFD), and Frontometaphyseal Dysplasia (FMD) and cardiospondylocarpofacial syndrome (CSCFS), respectively, rare multisystem syndromes, where CHD may appear in the clinical spectrum. We hypothesized that TAK1 plays a significant role in heart development and CHD and addressed this by genetic analysis in CHD patient cohorts and experiments in cell and animal models. Exome sequencing data from 1,471 CHD patients with extracardiac anomalies (syndromic CHD, sCHD), 2,405 patients with nonsyndromic CHD (nsCHD) and 45,082 controls showed increased burden of rare *TAB2* and *TAK1* variants in sCHD, but not in nsCHD. Detailed characterization of *tak1^-/-^*and *tab2^-/-^* zebrafish mutants revealed cardiac defects (dilated atrium, trabeculation defects, tachycardia and reduced contractility) as well as extracardiac developmental anomalies. RNA sequencing of *tak1^-/-^* mutant hearts showed downregulation of genes encoding core cardiac transcription factors, sarcomeric proteins and extracellular matrix proteins. Experiments with cell cultures and analysis of zebrafish larvae and gastruloids indicated that TAK1 via TAB2 and PKA-Cα is activated at the primary cilium during cardiomyogenesis and that TAK1 activation at this site is enhanced by cardiomyogenic signaling molecules, including ligands of the TGFB/BMP superfamily. Consistent with these findings, CRISPR/Cas9-mediated editing of TAK1 or administration of small molecule inhibitors targeting TAK1 inhibited ciliary signaling and cardiomyocyte differentiation *in vitro*, while FMD-causing mutations in TAK1 reduced its ciliary localization. In conclusion, our data establishes a central role for TAK1 and its upstream regulators in cardiac development and syndromic CHD, coordinated via the primary cilium.

## Introduction

The heart is the first organ to form during human embryonic development. Heart development is a complex process, orchestrated spatially and temporally by networks of morphogens, signaling pathways, transcription factors and chromatin modulators, which regulate proliferation, differentiation, and positioning of cardiac progenitor cells ^1,2^. A significant part of these processes have been linked to motile or primary cilia ^3,4^, which in the developing embryo project from non-proliferating cells and exchange signals with the surrounding tissue ^5,6^. However, coordination of cardiogenic signaling events by primary cilia remains poorly understood, posing a significant challenge in unraveling the underlying causes of Congenital Heart Disease (CHD), a variety of structural malformations of the heart or intra-thoracic vessels that affect cardiac function ^7,8^. Clinically, CHD may present as the only birth defect in the newborn (non-syndromic CHD, nsCHD) or together with extracardiac anomalies (syndromic CHD, sCHD).

Transforming Growth Factor-Beta-Activated Kinase 1 (TAK1, also known as MAP3K7) is part of the non-canonical branch of TGFB/BMP signaling and is activated by phosphorylation mediated by the catalytic alpha subunit of cAMP-dependent protein kinase (PKA-Cα, encoded by the *PRKACA* gene) and via interaction with TAK1 Binding Protein 2 (TAB2)^9,10^. Rare genomic and intragenic variants, which disrupt the normal function of *TAB2* have been associated with TAB2 haploinsufficiency syndrome (THS) ^11,12^. Rare gain-of-function or loss-of-function variants in *TAK1* can cause frontometaphyseal dysplasia (FMD, OMIM #617137) or Cardiospondylocarpofacial syndrome (CSCF, OMIM #157800) ^13,14^ and rare mutations in *PRKACA* give rise to cardioacrofacial dysplasia (CAFD, OMIM #619142) ^15–17^. These syndromes are all multisystem disorders, which appear to have both distinct and overlapping clinical characteristics, including cardiac defects. *TAB2* was initially identified as a disease gene via genetic analysis of CHD patients ^18^, and clinical analyses of THS patients show that CHD and cardiomyopathy are found in 86% and 51% of patients, respectively ^11^. A significant frequency of CHD has also been observed among patients with rare variants in *TAK1* and *PRKACA* ^17,19^. Based on the high frequency of cardiac defects in these patients, we hypothesized that TAK1, TAB2 and PKA-Cα might be integral components of a collaborative signaling axis that regulates developmental processes, prominently contributing to cardiogenesis.

The current study was undertaken to address this hypothesis and advance our understanding of CHD by exploring the role of TAK1 and signaling partners in cardiogenesis. Zebrafish *tak1* and *tab2* homozygous mutant larvae recapitulate several of the developmental defects observed in FMD/CSCF and THS patients, including heart malformations, and RNA sequencing of dissected hearts suggests that Tak1 functions upstream of core cardiac transcription factors and is involved in development of both extracellular matrix (ECM) and myocardium. By combining zebrafish investigations with analyses in gastruloids and cell model systems, we further provide evidence that TAK1 operates in concert with TAB2 and PKA-Cα in a non-canonical branch of TGFB/BMP signaling at the primary cilium to control cardiomyogenesis. Further, mutations in TAK1 and/or pharmacological suppression of its activity impede the activation of JNK1/2 at the cilium and disrupts the timely initiation and advancement of *in vitro* cardiomyocyte differentiation, underscoring the critical involvement of TAK1 in cilium-mediated cardiogenesis. These findings provide novel insight into heart development and how ciliary signaling defects converge on sCHD.

## Results

### Increased burden of rare *TAK1* and *TAB2* protein altering variants in syndromic CHD

To investigate the association between *TAK1*, *TAB2, PRKACA* and CHD, we initially analyzed exome sequencing (ES) data from a cohort of 1,471 sCHD patients, 2,405 nsCHD patients and 45,082 controls ^20^. We calculated the burden of rare (MAF<0.001) synonymous (SYN) and protein altering variants (PAVs) between the two patient cohorts and controls and analyzed the difference using Fisher’s exact test. In ES data obtained from sCHD patients we observed an approximately two-fold increase in the burden of rare PAVs for *TAK1, TAB2 and PRKACA*, when compared to controls. This difference was statistically significant for *TAK1* and *TAB2* (P-value 0.042 and 0.0081, respectively) (**Fig. 1a-c**). In ES data obtained from nsCHD, we observed no difference in variant burden for any of the three genes (not shown). As a negative control, we compared the burden of rare synonymous variants, and did not observe any significant difference between nsCHD and controls (**Fig. 1a-c**). Together, these results indicate that rare variants in *TAK1* and *TAB2* are associated with sCHD.

**Figure 1.**
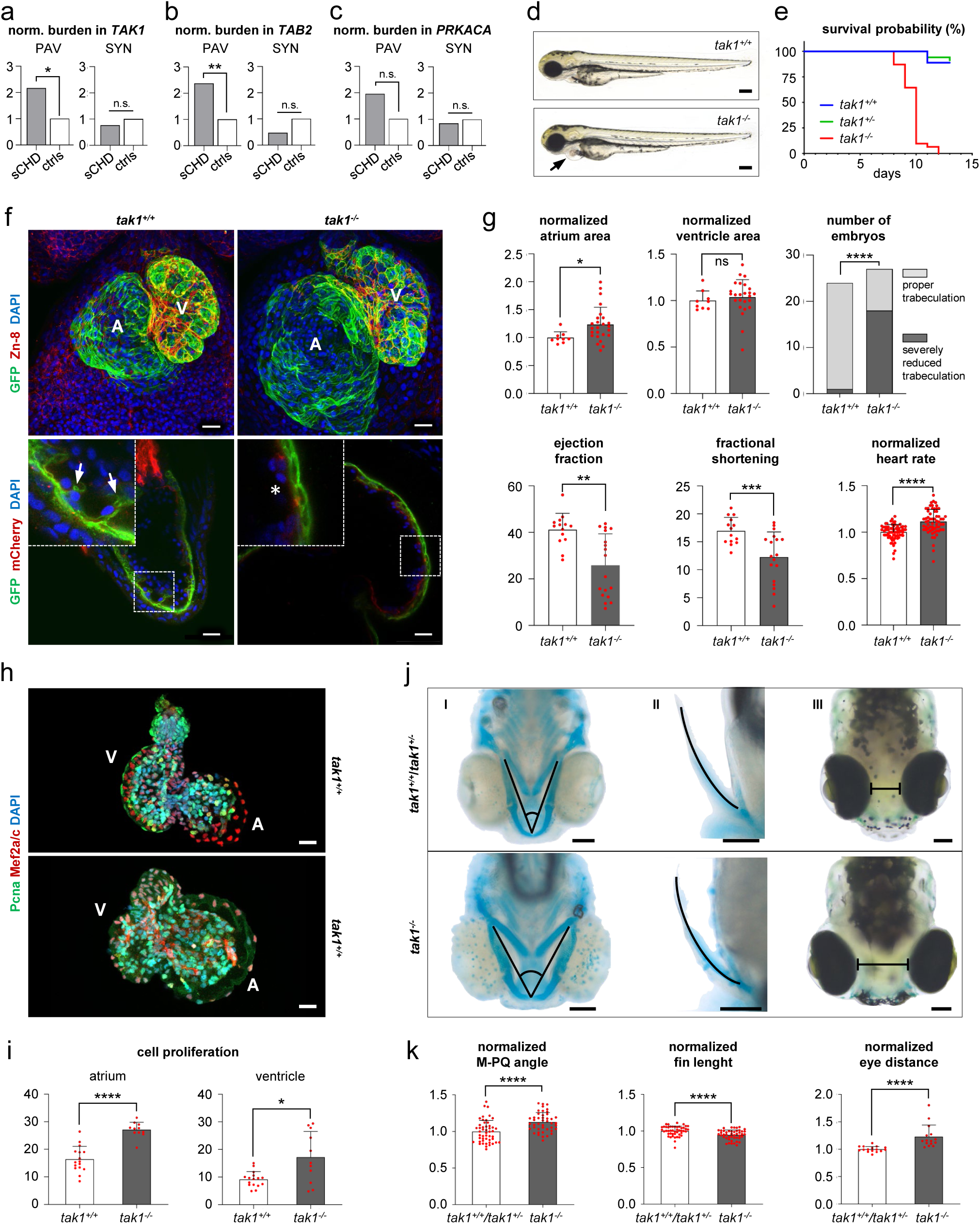
Phenotype of zebrafish *tak1* mutants. **a-c.** Normalized burden of rare protein altering variants, PAV (left) and synonymous variants, SYN (right) in *TAK1* (a), *TAB2* (b) and *PRKACA* (c). The rare variants (MAF <0.001) were identified by exome sequencing of 1,471 CHD patients with extracardiac anomalies (sCHD) and 45,082 controls (ctrls). *: P<0.05, **: P<0.01. **d.** Gross morphology of 3 dpf *tak1^+/+^* and *tak1^-/-^* mutant zebrafish larvae highlighting pericardial edema (arrow). Scale bars, 0.2 mm. **e.** Survival of *tak1^+/+^, tak1^+/-^* and *tak1^-/-^* zebrafish larvae. **f.** Upper panel: IFM of transgenic (*Tg[myl7:GFP*]) *tak1^+/+^*and *tak1^-/-^* zebrafish hearts with anti-GFP (green) and zn-8 (red) antibodies at 3 dpf. A: atrium, V: ventricle. Lower panel: IFM of (*Tg[myl7:GFP;kdrl:mCherry]*) *tak1^+/+^* and *tak1^-/-^* zebrafish hearts with anti-GFP (green) and anti-mCherry (red) antibodies at 6 dpf. Ventricular trabeculae are indicated with arrows and trabeculation defects with asterisks. Nuclei were stained with DAPI (blue). Scale bars, 20 μm. **g.** Quantification of normalized ventricle area, atrium area, ejection fraction (%) and fractional shortening (%) at 3 dpf, heart rate at 5 dpf and trabeculation defects at 6 dpf. **h.** IFM of cardiomyocyte nuclei (DAPI, blue) and proliferating cells, using anti-Mef2a/c (red) and anti-Pcna (green) antibodies, respectively in 3 dpf *tak1^+/+^*(upper) and *tak1^-/-^* (lower) zebrafish heart extracts. Scale bars, 20 μm. **i.** Quantification of cell proliferation index in atrium (left) and ventricle (right) presented in (h). **j.** Extracardiac defects in *tak1* mutants. Bright-field images of *tak1^+/+^/tak1^+/-^* (upper panel) and *tak1^-/-^* (lower panel) larvae for (I) Meckel’s-palatoquadrate (M-PQ) angle measurements of cartilage with Alcian blue staining at 5 dpf. Black lines show the measured angle in larvae. (II) Measurement of fin length with Alcian blue staining at 5 dpf. Black line shows measured fin length. (III) Eye distance measurement at 7 dpf. Black line shows the measured eye distance in larvae. Scale bars, 0.1 mm. **k.** Quantification of measurements obtained from panels I-III in (j). *: P<0.05, **: P<0.01, ***: P<0.001, ****: P<0.0001, ns: not significant.

### Homozygous mutation of *tak1* and *tab2* is lethal and cause heart defects

To analyze the involvement of TAK1 in *in vivo* heart development, we generated zebrafish lines with 1 bp and 8 bp deletion in the coding sequence of *tak1* and *tab2*, respectively, using the CRISPR-Cas9 technique (**Suppl. Fig. 1 a-c**). Both variants lead to a shift in the reading frame of the gene, resulting in a premature stop codon.

Zebrafish *tak1^-/-^* homozygote mutants (will be referred to as *tak1* mutants hereafter) displayed apparent pericardial edema by 3 days post fertilization (dpf) with a hundred percent penetrance while the heterozygote siblings were phenotypically inseparable from the wildtype siblings (**Fig. 1d**). Survival of *tak1* mutants was severely compromised compared to wildtypes and heterozygotes, with no *tak1* mutant surviving past 12 dpf (**Fig. 1e**). Immunofluorescence microscopy (IFM) analysis of the cardiomyocytes revealed defects in cardiac morphology. We observed large variability in ventricle and atrium size in *tak1* mutants, with the atrium being significantly enlarged at 3 dpf (**Fig. 1f, g**). When we assessed the hearts at 6 dpf, we observed significantly reduced trabeculation in *tak1* mutants (**Fig. 1f, g**). The *tak1* mutants had defects in heart function, as measured by reduced fractional shortening and ejection fraction at 3 dpf, and in addition, we observed increased heart rate in *tak1* mutants at 5 dpf (**Fig. 1g**).

To address the changes in the cardiac chamber sizes in *tak1* mutants, we counted their atrial and ventricular cell numbers, measured the size of the atrial and ventricular cardiomyocytes, and compared them to those of *tak1^+/+^*larvae. We observed significantly more cells in the atrium of *tak1* mutants (**Suppl. Fig. 2a, b**) but did not observe any difference in cell size (**Suppl. Fig. 2c, d**). We observed a significant increase in cardiomyocyte proliferation index both within the atrium and ventricle of *tak1* mutants at 3 dpf (**Fig. 1h, i**). To assess gross effects on myofibrils, we measured the distance between sarcomeric z-discs in atrium and ventricle but did not identify any difference between *tak1* mutants and wildtype larvae (**Suppl. Fig. 2e, f**). Cardiac valve defects are hallmarks of THF and are also found in a number of patients with *TAK1* mutation ^11,13^. Thus, we assessed the atrioventricular (AV) valves in *tak1^-/-^* (and *tab2^-/-^*, see below) mutants. Although, some of the *tak1* mutants exhibited misnumbered cells in the valves, the overall number of cells were not significantly altered (**Suppl. Fig. 2g, h**). Finally, we tested the *tak1* mutants for increased apoptosis by acridine orange staining and analysis of expression of *bcl2* and *baxa*, but there was no difference in apoptosis markers between *tak1* mutant and *tak1^+/+^* larvae **(Suppl. Fig. 2i, j**).

To further analyze the effect of defective Tak1 signaling in heart development, we created *tab2^-/-^* homozygous mutants (will be referred to as *tab2* mutants hereafter). These mutants phenocopied *tak1* mutants with a slight delay. The *tab2* mutants started displaying apparent pericardial edema at 3 dpf, which became hundred percent penetrant by 5 dpf. In addition, we noted that *tab2* mutants, similar to *tak1* mutants at 5 dpf, display a protruding mouth (**Suppl. Fig. 3a**). We observed less pronounced lethality in *tab2* mutants, beginning from 7 dpf, with survival probability of 40% at 19 dpf (**Suppl. Fig. 3b**). However, *tab2* mutants did not survive to adulthood, suggesting that significant mortality of mutants occurs between 20 dpf and adulthood. Similar to *tak1* mutants, cardiac morphological defects in chamber size, looping and trabeculation were also observed in the *tab2* mutants; which displayed enlarged atria at 5 dpf, with severely reduced ventricular trabeculation observed at 6 dpf (**Suppl. Fig. 3c**). Cardiac function was also compromised in *tab2* mutants, marked by increased heart rate and reduced contractility at 5 dpf (**Suppl. Fig. 3d**). Cardiac valve defects are found in >75% of patients with *TAB2* mutations ^11^. Consistently, *tab2* zebrafish mutants displayed AV valve developmental defects, marked with a significant decrease in cell number (**Suppl. Fig. S3e, f**).

Finally, to validate the cardiac phenotypes in mutant larvae, we injected wildtype fertilized oocytes with antisense morpholino oligonucleotide (MO) targeting *tak1* and *tab2*, respectively. The cardiac phenotypes we observed in the morphants were similar to those of the *tak1* and *tab2* mutants, and could be rescued by co-injection with wildtype RNA (**Suppl. Fig. S4a, b**). Thus, the MO experiments support the genotype-phenotype correlation in mutants. In addition, co-injection experiments suggested a genetic interaction between *tak1* and *tab2* during heart development (**Suppl. Fig. 4c, d**).

### *tak1* and *tab2* mutants exhibit abnormalities in both craniofacial structure and fin development

Facial dysmorphisms and limb defects are among the clinical characteristics of FMD, CSCF and THS ^11–14^. To test if *tak1* and *tab2* mutants phenocopy the human syndromes, we compared craniofacial features and length of the pectoral fins in homozygous mutants and wildtype/heterozygous siblings. Homozygous *tak1* and *tab2* mutants showed significant increase in Meckel’s-Palatoquadrate angle at 5 dpf. Eye defects were found in *tab2* mutants at 5 dpf and in *tak1* mutants at 7 dpf, respectively. Altogether, these data suggest craniofacial abnormalities in the mutants (**Fig. 1j, k**; **Suppl. Fig. 3g, h**). At 5 dpf, the fin lengths of both *tak1* and *tab2* mutants were significantly decreased, suggesting brachydactyly. We did not observe significant phenotypes in adult heterozygous *tak1^+/-^* and *tab2^+/-^* and the lethality of homozygous genotypes prevented us from analyzing adult *tak1* and *tab2* mutants. Together with the observed cardiac defects, these results suggest that the *tak1* and *tab2* mutants phenocopy several of the clinical characteristics of human FMD, CSCF and THS.

### Transcriptome analysis show downregulation of genes encoding core cardiac transcription factors, sarcomeric proteins and extracellular matrix proteins in *tak1* mutants

To identify developmental mechanisms regulated by TAK1, we compared the cardiac transcriptome of wildtype and mutant *tak1* larvae. To this end, we dissected the hearts of transgenic (*Tg[myl7:GFP*]) larvae at 3 dpf and analyzed the transcriptome of the hearts by RNA sequencing. We identified 581 downregulated and 863 upregulated genes (**Fig. 2a**). These genes correspond to 474 and 712 human orthologues, respectively (listed in **Suppl. Table 1**). Within the list of downregulated genes, we observed enrichment of genes known to cause CHD in human patients and mouse models ^21,22^, suggesting that TAK1 promotes the expression of several genes known to cause CHD (**Suppl. Fig. 5a**). We observed no significant enrichment of known CHD genes within the list of upregulated genes. Gene-ontology (GO) analysis of the human orthologues to downregulated genes, showed enrichment of genes involved in heart morphogenesis, cardiomyogenesis, muscle tissue development and ECM (**Fig. 2b, left panel**). Significantly downregulated genes include genes encoding cardiac transcription factors and sarcomeric proteins as well as genes involved in structure and metabolism of the ECM (**Fig. 2c**). Downregulation of the cardiac transcription factor genes *tbx5b*, *gata4* and *hand2*, sarcomeric genes *actc1a* and *myl7*, and ECM related gene *has2* in *tak1* mutants was confirmed by RT-qPCR analysis of dissected hearts from 3 dpf larvae (**Suppl. Fig. 5b**). Comparison of the cardiac expression level of the same set of genes between *tab2^+/+^*and *tab2* mutant larvae at 5 dpf gave similar results (**Suppl. Fig. 5c**), supporting functional interaction between *tak1* and *tab2* in heart development. GO analysis of upregulated genes suggested that these genes were involved in basic cellular processes, differentiation of myeloid cells and platelet degranulation (**Fig. 2b, right panel**), thus a significant part of the upregulated genes may be false positives, caused by entrapment of more blood cells in the dilated atrium of mutant hearts, compared to wildtype hearts.

**Figure 2.**
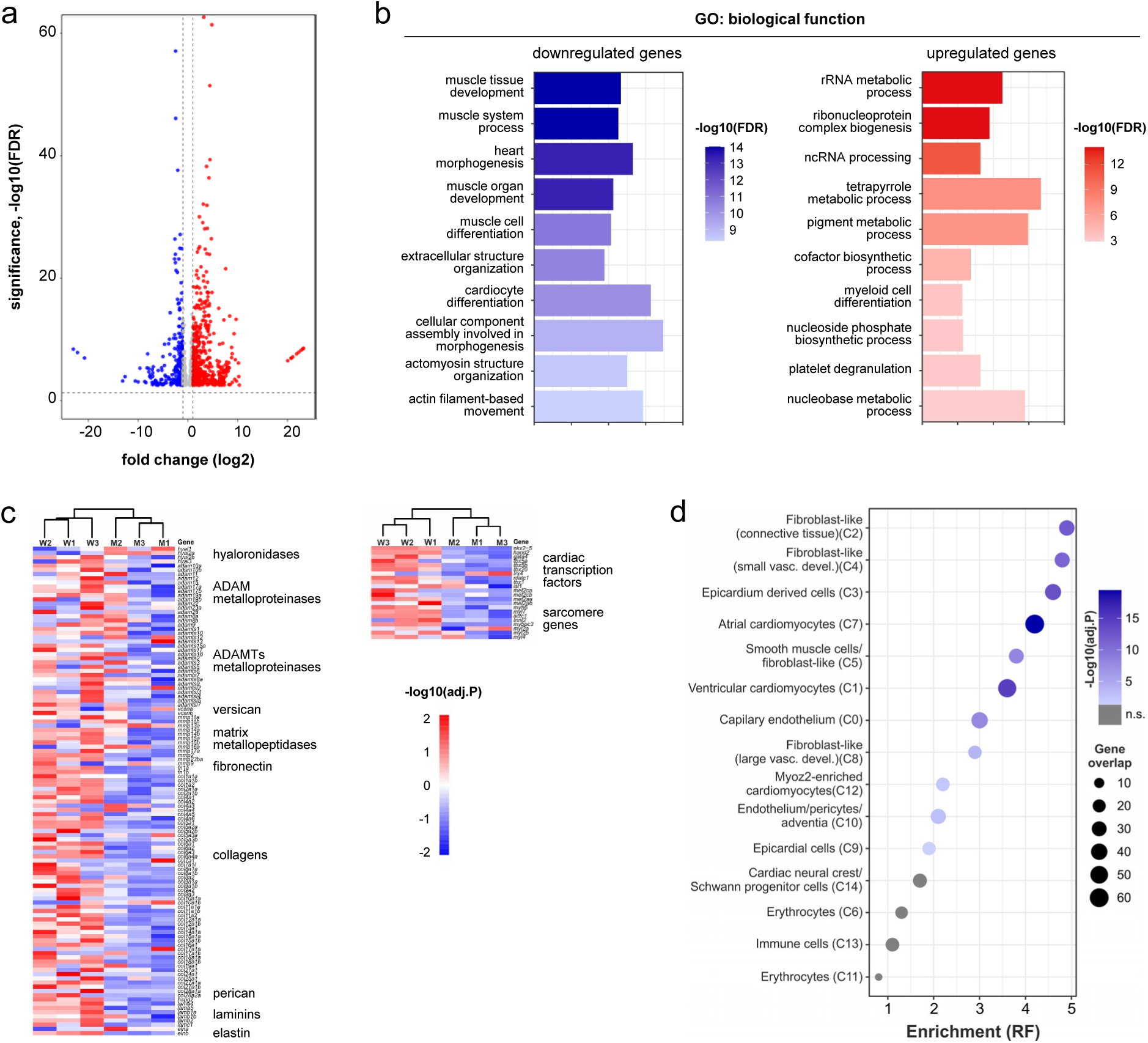
Transcriptome analysis of dissected zebrafish hearts. RNA sequencing was performed on hearts dissected from 3 dpf *tak1^+/+^* and *tak1^-/-^*zebrafish larvae. **a.** Volcano plot of 581 down (blue) and 863 upregulated (red) genes. **b.** Gene ontology (GO) enrichment analysis of 474 down (left) and 712 upregulated (right) genes (human orthologues to zebrafish genes). **c.** Heatmap showing the expression values of selected genes associated with ECM (left) and cardiomyogenesis (right). **d.** Enrichment of downregulated genes in the expression signature of fifteen cell-types in 6.5-7 WPC human developing hearts ^23^. Enrichment is shown as representation factor (RF). A hypergeometric distribution was used to test the significance.

Lastly, to identify possible cardiac cell types affected by downregulated genes, we calculated the enrichment of downregulated genes in cell-specific gene-signatures obtained from single cell sequencing of human embryonic hearts ^23^. We observed significant enrichment of downregulated genes in most cell types of the developing heart, except neural crest cells (**Fig. 2d**). Most pronounced enrichment was observed in cardiac fibroblasts, epicardium derived cells, cardiomyocytes and smooth muscle cells. For upregulated genes, we only observed enrichment in gene-sets specific for erythrocytes and immune cells, supporting that a significant part of upregulated genes is related to blood cells (**Suppl. Fig. 5d**).

Based on the cardiac transcriptome analysis, we conclude that TAK1 coordinates a significant part of gene-networks involved in development of the myocardium, cardiac ECM and cardiac smooth muscle cells.

### TAK1 localizes to primary cilia and is activated at this site by TGFB/BMP stimulation and during cardiomyogenesis

In the developing heart, canonical TGFB/BMP signaling is coordinated by the primary cilium ^24,25^. To investigate whether TAK1 similarly functions at the primary cilium during cardiogenesis, we initially analyzed the subcellular localization of Tak1 in zebrafish hearts at 3 dpf. Whole-mount IFM analysis of dissected wildtype hearts showed that Tak1 predominantly concentrates to the basal region of primary cilia in cardiomyocytes in the atrium, ventricle, and outflow tract (**Figure 3a, left panel**), whereas immunoreactivity is absent in hearts dissected from *tak1* mutants (**Figure 3a, right panel**), validating antibody specificity and ciliary Tak1 localization.

**Figure 3.**
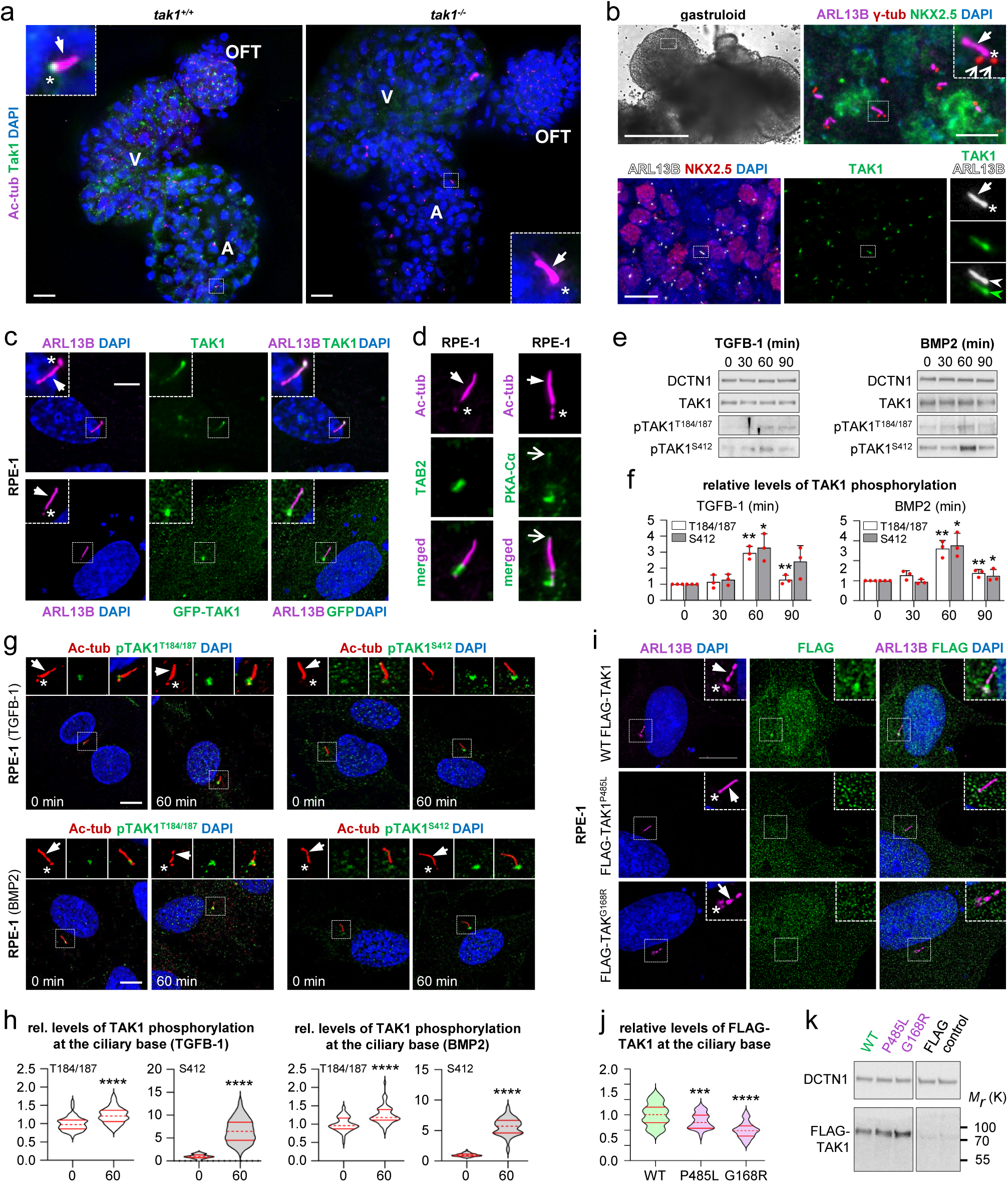
TAK1 localizes to the primary cilium and is activated at this site by TGFB/BMP stimulation. **a.** IFM of extracted *tak^+/+^* (left) and *tak1^-/-^*(right) hearts stained with Tak1 (green) and acetylated tubulin antibody (Ac-tub) (pseudocoloured magenta) at 3 dpf. A: atrium, V: ventricle, OFT: outflow tract. Scale bars, 20 μm. **b.** Upper left panel: DIC microscopy image of day 7 gastruloid, white box indicates ROI. Scale bar, 0.5 mm. Upper right panel: presence of primary cilia (ARL13B, magenta) in cardiac progenitor cells (NKX2.5, green). Centrioles were stained with γ-tubulin (red). Scale bars, 10 μm. Lower row panels: NKX2.5 (green) positive cardiac progenitor c ells and ARL13B (pseudocoloured white) stained primary cilia. TAK1 (green) is observed at primary cilia of day 7 gastruloids. Scale bars, 10 μm. **c.** IFM of endogenous TAK1 (green) and GFP-tagged TAK1 (green) at the primary cilium (ARL13B, pseudocolored magenta) in RPE-1 cells. Scale bar, 5 μm **d.** IFM images of TAB2 and PKA-Cα, (green) at the primary cilium (Ac-tub, pseudocoloured magenta). **e.** Representative images of WB analysis of TAK1, pTAK1^T184/187^ and pTAK1^S412^ at indicated time points in RPE-1 cells upon TGFB-1 (left) and BMP2 (right) stimulation. DCTN1 was used as a loading control. *: P<0.05, **: P<0.01 **f.** Quantification of relative levels of TAK1 phosphorylation with TGFB-1 (left) and BMP2 (right) ligand stimulation from (e). **g.** Representative IFM images of pTAK1^T184/187^ and pTAK1^S412^ (green) at time point 0 and 60 minutes upon TGFB-1 (upper panels) and BMP2 (lower panels) stimulation in RPE-1 cells. Cilium is marked with Ac-tub (pseudocolored magenta). Scale bars, 10 μm. **h.** Quantification of relative levels of fluorescent intensities of pTAK1 proteins at the ciliary base upon ligand stimulation from (g). **i.** Representative IFM images of FLAG-tagged WT TAK1 and TAK1 FMD patient variants (green) costained with ARL13B (pseudocolored magenta) in transfected RPE-1 cells. Scale bar, 10 μm. **j.** Quantification of relative levels of FLAG-tagged WT TAK1 and mutant TAK1 at the ciliary base from (i). ***: P<0.001, ****: P<0.0001. **k.** WB analysis showing expression levels of FLAG-tagged WT TAK1 and TAK1 FMD patient variants in transfected RPE-1 cells. In IFM images, arrows and asterisks indicate ciliary axoneme and base, respectively.

To further substantiate ciliary localization of TAK1 during cardiomyogenesis, we performed IFM analyses in day 7 gastruloids (**Fig. 3b, upper left panel**), which differentiate into several progenitor cell populations, including heart field cells, in a 3D manner (**Suppl. Fig. 6a**) ^26^. Our investigation revealed distinct regions within the gastruloids characterized by an abundance of the cardiac transcription factor, NKX2-5, and the first heart field marker HCN4 ^27^ (**Suppl. Fig. 6b**), demonstrating the presence of cardiac progenitors ^27,28^. Further investigation highlighted that NKX2-5-positive regions developed distinct Troponin-I cardiomyofibrils (**Suppl. Fig. 6c**). Co-staining of NKX2-5 and HCN4 was evident in numerous regions throughout the gastruloids and we found that cardiac progenitor cells expressing either NKX2-5 or both NKX2-5 and HCN4 form primary cilia (**Fig. 3b, upper right panel**), which serve as the principal site for the localization of TAK1 (**Fig. 3b, lower row**). These discoveries align with our observations in the developing heart of zebrafish larvae, pinpointing the presence of TAK1 in primary cilia during cardiogenesis. Furthermore, these novel findings underscore the utility of gastruloids as a system through which the involvement of primary cilia in early mammalian development can be delineated.

To support these findings and examine ciliary TAK1 activity in the context of cardiomyogenic stimuli, we proceeded to investigate TGFB/BMP-mediated TAK1 activation in retinal pigment epithelial (RPE-1) cells, which are well-established model cells for developmental signaling and primary cilia studies ^5^. In pursuit of this objective, we initially conducted IFM analysis, uncovering significant accumulation of both endogenous and GFP-tagged TAK1 at the ciliary base, and in some cases with notable distribution along the primary cilium (**Fig. 3c**). Furthermore, we observed prominent ciliary base localization of the essential upstream regulators of TAK1 activity, TAB2 and PKA-Cα, which also localized to the ciliary tip (**Fig. 3d**). Localization of PKA-Cα to primary cilia aligns with previous findings on the role of this subunit in the regulation of Hedgehog (Hh) signaling in the cilium, where it phosphorylates and inactivates GLI2/3 transcription factors to repress expression of Hh target genes ^6,29^. Further, TAK1 and TAB2 were recently identified in cilia proteomic studies ^30,31^.

Next, western blot and IFM analyses were carried out to evaluate TGFB/BMP-mediated phosphorylation of TAK1 at T184/187 and S412, which marks its activation mediated by TAB2 and PKA-Cα respectively ^10,32^. To this end, cells were subjected to stimulation over a time interval of 90 mins with TGFB-1 and BMP2, which play crucial roles in cardiac development, including, but not limited to, intricate processes of cardiomyogenesis ^24^. Both ligands elicited phosphorylation of TAK1 at T184/187 and S412, reaching peak levels at 60 minutes of stimulation (**Fig. 3e, f**), and at this time point, increased phosphorylation levels were observed specifically at the base of the primary cilium (**Fig. 3g, h**). Because TGFB/BMP receptors function at the level of primary cilia ^5,24^, these results support the conclusion that cardiomyogenic signaling molecules of the TGFB/BMP family operate through cilia-localized TAB2 and PKA-Cα to activate TAK1 at this site.

As individuals with TAK1 mutations may exhibit the multi-system syndrome FMD ^14^, we investigated the potential impact of patient-specific mutant forms of TAK1 on its localization to primary cilia by transiently transfecting RPE-1 cells with plasmids coding for wildtype FLAG-TAK1 or plasmids coding for patient-specific FLAG-TAK1 variants G168R and P485L ^14^. IFM analysis of transfected cells validated the ciliary localization of wildtype TAK1, whereas the patient-specific variants showed impaired recruitment to this site (**Fig. 3i-k**). Thus, TAK1 mutations identified in FMD patients appear to influence the localization of TAK1 to the primary cilium, potentially compromising cardiogenesis through impaired ciliary signaling.

### TAK1 is required for *in vitro* cardiomyocyte differentiation

To corroborate our findings in zebrafish, gastruloids, and RPE-1 cells, and to delve deeper into the role of TAK1 in cardiomyocyte differentiation, we finally opted for P19CL6 teratocarcinoma-derived pluripotent stem cells, which serve as a suitable model for studying early cardiomyogenesis ^33–36^. P19CL6 cells exit their pluripotent stage within a few days of DMSO treatment, accompanied by the initiation of cardiomyogenesis (**Fig. 4a**), as indicated by a decrease in SOX2 expression and an increase in the expression of transcription factor GATA4, which promotes cardiac development and differentiation of the myocardium (**Fig. 4b**). Within 12 days of treatment, the cells form functional clusters of beating cardiomyocytes, displaying well-defined striated sarcomeric patterns of Troponin I and ⍺-actinin (**Figure 4a-d**), and throughout their differentiation the cells display primary cilia (**Fig. 4e**), which are required for cardiomyogenesis in this stem cell model ^24,33^.

**Figure 4.**
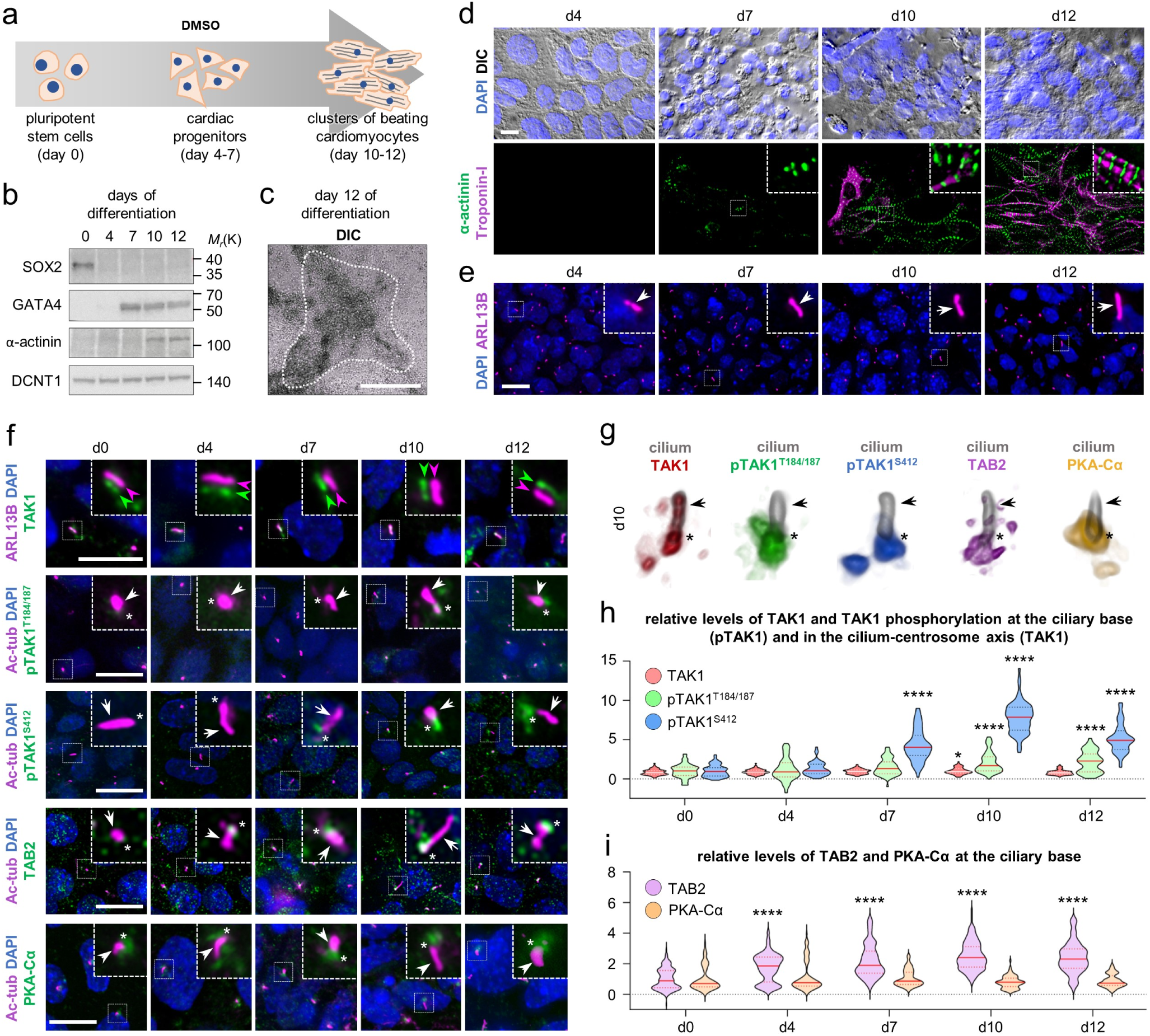
TAK1 is activated at the primary cilium during *in vitro* cardiomyocyte differentiation. **a.** Graphical illustration of cardiomyocyte differentiation from day 0 to day 12 upon stimulation of pluripotent stem cells (P19CL6) with 1% DMSO. b. WB analysis of SOX2, GATA4 and α-actinin protein levels in P19CL6 cells upon DMSO stimulation from day 0 to day 12. DCTN1 was used as a loading control. c. DIC of a beating cluster of cardiomyocytes (within dotted white lines) at day 12 of DMSO stimulated P19CL6 cells. Scale bar, 1 mm. d. DIC and IFM images showing P19CL6 cells undergoing differentiation from day 4 to day 12. Upper panels: differential interference contrast (DIC) microscopy with staining of nuclei (DAPI, blue). Lower panels: IFM images showing ⍺-actinin (green) and Troponin-I (pseudocolored magenta) in the same area as the upper panels. Scale bar, 10 µm. e. Representative IFM images displaying primary cilia (arrows, ARL13B, pseudocolored magenta) during cardiomyocyte differentiation from day 4 to day 12. Scale bar, 10 μm. f. Representative IFM images showing expression of TAK1 (shifted), pTAK1^T184/187^, pTAK1^S412^ and upstream components TAB2 and PKA-Cα (all in green), co-stained with ciliary markers (ARL13B or Ac-tub, pseudocolored magenta) in P19CL6 cells. Arrow marks the cilium and asterisks denotes the ciliary base region. Scale bar, 20 μm. g. Voxel view of primary cilia (pseudocolored grey) along with TAK1, pTAK1^T184/187^, pTAK1^S412^, TAB2 and PKA-Ca, pseudocolored as red, green, blue, magenta, and yellow, respectively, at day 10 of P19CL6 cell differentiation. h, i. Quantification of relative levels of TAK1 and activated TAK1 proteins (h) and TAB2, PKA-Cα (i) in the cilium-centrosome axis and at the ciliary base from (f). Statistical analysis was performed via Kruskal-Wallis test for TAK1 proteins and PKA-Cα, one-way ANOVA was performed for TAB2, ****: P<0.0001.

During P19CL6 cardiomyogenesis, we observed a substantial enrichment of TAB2 as well as phosphorylation of TAK1 at T184/187 and S412 at the primary cilium, reaching peak levels around day 10 of differentiation (**Fig. 4f-i**). In contrast, the levels of total TAK1 and PKA-Cα at the cilium remained relatively stable throughout the differentiation process (**Fig. 4f-i**). Highlighting day 10 of differentiation, 3D imaging further delineated localization of TAK1 around the ciliary base as well as along the entire length of the cilium in a punctate pattern, whereas PKA-Cα, TAB2 and phosphorylated versions of TAK1 predominantly localized to the ciliary base region (**Fig. 4g**). Together these findings suggest that TAB2 is recruited to the cilium during differentiation, where, in conjunction with PKA-Cα, TAK1 phosphorylation and activation is mediated in the context of cardiomyogenesis.

To examine the necessity of TAK1 for cardiomyogenesis, we next generated CRISPR-Cas9 P19CL6 mutants of *Tak1* containing a deletion, P19CL6*^Tak1Δ^*^85^*^/Δ^*^85^, resulting in skipping of exon 2, thus causing in-frame deletion of amino-acid residue 41-77 within the kinase domain of TAK1 (**Suppl. Fig. 7a, c**). The mutant cells exhibited prolonged expression of SOX2 and a delayed onset of GATA4 expression compared to wildtype (WT) clone cells (P19CL6^WT^) during the DMSO stimulation period (**Fig. 5a-d**). At day 7, while the majority P19CL6^WT^ cells were GATA4 positive, most P19CL6*^Tak1Δ^*^85^*^/Δ^*^85^ remained OCT3/4-positive and GATA4-negative (**Fig. 5e, f**). Additionally, *Tak1* mutants displayed significant deficiencies in the formation of striated ⍺-actinin structures, noticeably reduced by day 12 of DMSO treatment, accompanied by little or no expression of Troponin-I (**Fig. 5e, g**). Furthermore, we generated CRISPR-Cas9 P19CL6 clones with a knockout of Tab2, P19CL6*^Tab^*^2^*^-/-^* (**Suppl. Fig. 7b, c**). Similar to TAK1 mutant cells, P19CL6*^Tab^*^2^*^-/-^* cells exhibited severe delay in cardiomyogenesis, as indicated by reduced expression of GATA4 and the absence of striated sarcomeric patterns of Troponin I and ⍺-actinin (**Suppl. Fig. 7d-h**). Hence, mutations in *Tak1* and *Tab2* hinder cardiomyocyte differentiation.

**Figure 5.**
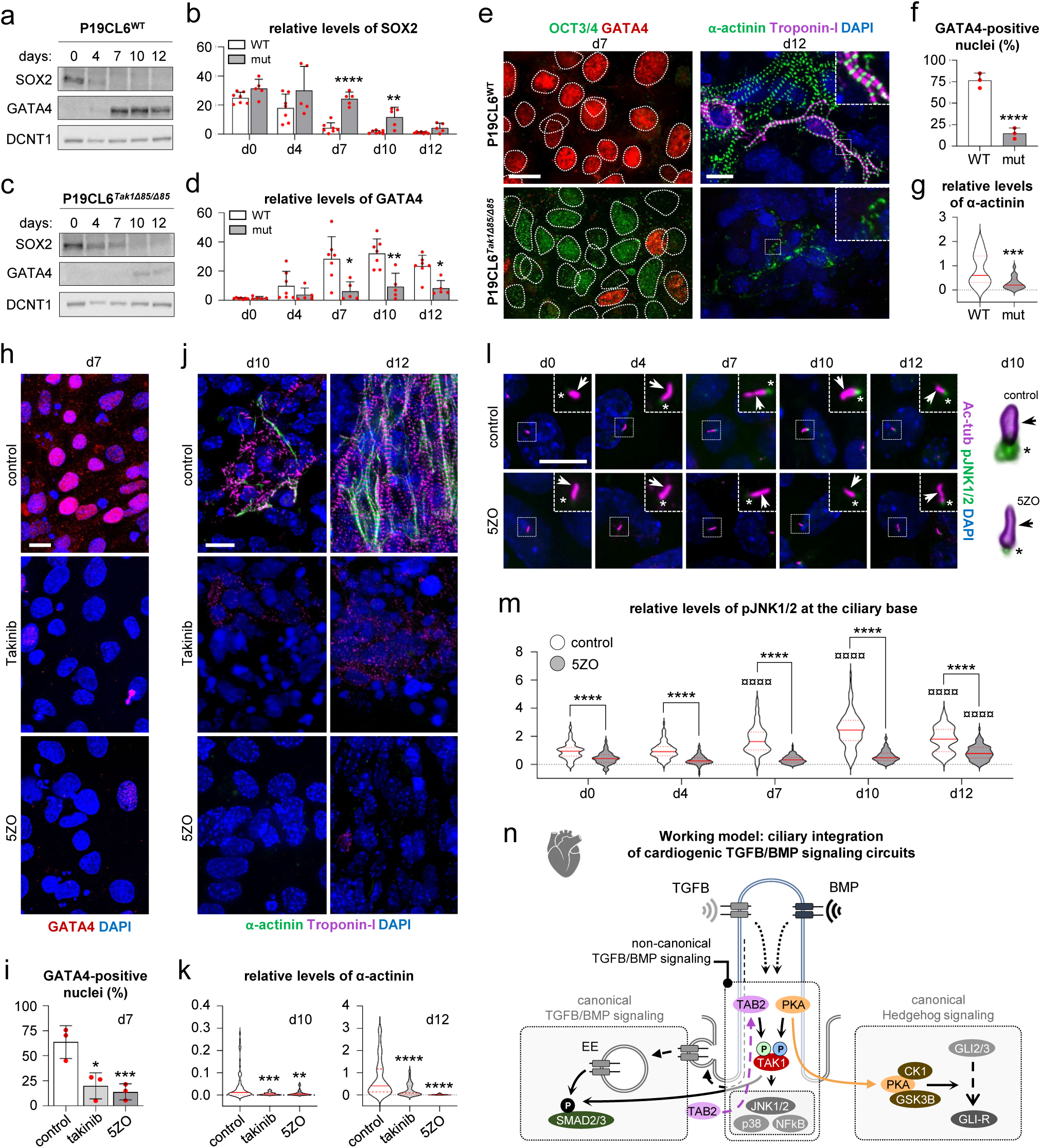
Mutation of TAK1 or inhibition of TAK1 activity impairs *in vitro* cardiomyocyte differentiation. **a, c.** WB analysis at day 0, 4, 7, 10 and 12 of differentiation using antibodies against SOX2 and GATA4 from DMSO treated P19CL6^WT^ and P19CL6*^Tak1Δ^*^85*/*^ *^Δ^*^85^ cells, respectively. DCTN1 was used as a loading control. **b, d.** Quantification of relative levels of SOX2 and GATA4 protein from a and c. **e.** Representative IFM images of OCT3/4 (green) and GATA4 (red) localization at day 7 (left panels) and α-actinin, troponin-I at day 12 (right panels) in DMSO induced P19CL6^WT^, P19CL6 Tak1^KO^ cells. Scale bars, 10 µm. **f.** Percentage of GATA4 positive nuclei from e (left panel). **g.** Quantification of relative levels of α-actinin from day 12 from e (right panel). **h, j.** Representative IFM images of GATA4 (red) at day 7 of differentiation (h) and α-actinin (green) and Troponin-I (pseudocolored magenta) at day 10 and 12 of differentiation (j) from P19CL6^WT^ cells treated without (control) and with TAK1 inhibitors (Takinib and 5ZO). Scale bars, 10 µm. **i, k.** Percentage of GATA4 positive nuclei from (h) and relative levels of α-actinin from (j). **l.** Representative IFM images of pJNK1/2 (green) at the primary cilium (pseudocolored magenta) from control and 5ZO-treated DMSO induced P19CL6^WT^ cells from day 0 to day 12. Scale bar, 10 μm. Right panel: Representative voxel view of pJNK1/2 at the primary cilium at day 10 of P19CL6 differentiated cells in control and 5ZO-treated cells. **m**. Relative levels of pJNK1/2 at the ciliary base from (l). **n.** Working model for TAB2 and PKA-mediated activation of TAK1 and its downstream signaling components at the primary cilium to induce cardiogenesis. ****: P<0.0001. *: P<0.05, **: P<0.01, ***: P<0.001, ****: P<0.0001. ¤¤¤¤: P<0.0001 compared to the respective means at day 0.

To confirm the results for *Tak1* mutant cells, we exposed P19CL6 cells to TAK1 inhibitors, Takinib and (5Z)-7-oxozeanol (5ZO)^37,38^ throughout the differentiation protocol. In the presence of either inhibitor, P19CL6 cells displayed a significant reduction in GATA4 positive nuclei by day 7 of DMSO stimulation (**Fig. 5h, i**) as well as a marked reduction in formation of well-defined striated sarcomeric patterns of Troponin I and ⍺-actinin as compared to vehicle controls, with the most prominent inhibition of cardiomyocyte formation observed with 5ZO (**Figure 5j, k**). These studies confirm the CRISPR-Cas9 mediated mutant results, highlighting the importance of TAK1 in cardiomyogenesis.

TAK1 is acknowledged for its role in activating c-Jun N-terminal protein Kinase (JNK), p38 mitogen-activated protein kinase (p38), and Nuclear Factor kappa-light-chain-enhancer of activated B cells (NF-κB), all of which, through diverse mechanisms, play pivotal roles in regulating cardiac cell fate specification and the maturation of cardiomyocytes ^39,40^. To explore ciliary TAK1 activity in relation to downstream signaling events during cardiomyogenesis, we investigated the impact of TAK1 inhibition by 5ZO on the phosphorylation of JNK1/2 through IFM analysis in P19CL6 cells during the differentiation process. In control cells, we observed ciliary base localization of phosphorylated JNK1/2, which peaked at day 10 of DMSO treatment (**Figure 5l, m**). In contrast, the level of JNK1/2 phosphorylation remained low at the ciliary base throughout the experiment in the presence of 5ZO, and the levels were significantly lower than those observed in control cells (**Figure 5l, m**). These findings support the conclusion that the primary cilium serves as a critical hub for TAB2- and PKA-Cα-mediated activation of TAK1 in the non-canonical branch of TGFB/BMP signaling as well as for translating activation of this branch into downstream signaling events in cardiogenesis, which is presented by a working model in **Figure 5n**.

## Discussion

Variants in *TAK1*, *TAB2* and *PRKACA* are associated with rare human multisystem disorders, which may include CHD and cardiomyopathy as part of the clinical spectrum ^11–14,17,18^. Our analysis of exome sequencing data from a large CHD cohort, combined with *in vivo* and *in vitro* experiments support a significant role of both TAK1 and TAB2 in heart development and CHD. Importantly, we show that TAK1 operates in conjunction with TAB2 and PKA-Cα in non-canonical TGFB/BMP signaling at the primary cilium, and that FMD-associated mutations in TAK1 diminish its ciliary localization. These findings add to the growing evidence that underscores a significant role of cilia not only in cardiac left-right patterning but also in broader aspects of cardiac development and CHD ^3,4^.

In zebrafish *tak1* mutants we observed a significant increase in cell proliferation within the myocardium, compared to WT larvae, and cardiac transcriptome analysis showed that genes encoding core cardiac transcription factors and sarcomere proteins were significantly downregulated in *tak1* mutants. Supporting the role of TAK1 in cardiomyogenesis, mutation of *Tak1* or administration of small molecule inhibitors targeting TAK1 inhibited cardiomyocyte differentiation *in vitro*. In control cells, TAK1 function involved TAB2 recruitment, TAK1 phosphorylation and subsequent activation of downstream signaling via JNK1/2 at the primary cilium. Further substantiating a function of this signaling axis in cardiomyogenesis, *in vitro* cardiomyocyte differentiation is similarly obstructed in cells with knockout of TAB2. Based on these findings, we propose a model where the cilium takes center stage in coordination of cardiogenic signaling events where TAK1 in concert with other signaling pathways controls intrinsic processes of cardiogenesis. This signaling crosstalk include, but may not be limited to, the canonical branch of TGFB/BMP signaling and Hh signaling (**Fig 5n**). In support of this model, which emphasizes the intricate interplay among ciliary pathway modules in regulating developmental signaling outcomes, TAK1 has been shown to bind and regulate the activity of R-SMADs ^41^, and upstream of TAK1, PKA-Cα downregulates Hh signaling by processing GLI2/3 transcription factors within the ciliary compartment ^29^. Conversely, inhibiting GLI1 in the Hh pathway was shown to activate the TAK1-JNK1/2 signaling axis ^42^, introducing another layer of interaction between Hh and TAK1 signaling systems. We have also noted that TAK1 activation in a cell-type specific context is regulated by WNT5A ^41,43^, which modulates the length of primary cilia ^44^ and was proposed to operate via a ciliary TMEM67-ROR2 receptor complex in the planar cell polarity branch of WNT signaling ^45^. Our discovery of TAK1 signaling in the primary cilium thus paves the way for future studies to unravel the complexities of developmental signaling networks and how the coordination of such networks may operate in a temporal manner to control cardiogenic events.

In addition to its role in cardiomyogenesis, our cardiac transcriptome analysis of zebrafish larvae indicates that TAK1 regulates a broad range of genes related to the structure and metabolism of the cardiac ECM. Precise orchestration of ECM synthesis is crucial for establishment of the endocardial cushions (EC), the anlage of cardiac valves, which are cellularized by endothelial-to-mesenchymal transition (endoMT) of endocardial cells lining the EC ^46^. Thus, TAK1 dysregulation of cardiac ECM is likely to explain part of the valvular phenotypes, which are observed in patients with *TAK1* and *TAB2* variants ^11,19^. In support of this conclusion, TGFB/BMP signaling events critically regulate production and turnover of ECM across different contexts ^47^, and we speculate such events to converge on TAK1, influencing cardiac ECM formation and, consequently, establishment of endocardial cushions.

Moreover, ECM integrity impacts the bioavailability of TGFB/BMP ligands ^47^, and primary cilia themselves interact with ECM ^48^. However, further investigations are required to fully understand how dysregulation of TAK1 manifest in valvular phenotypes within a ciliary context. Our observation of a reduced number of Sox9-expressing mesenchymal cells in zebrafish *tab2* mutants, but not in *tak1* mutants, also suggests that TAB2 might play a TAK1-independent role in regulation of endoMT. Such a role is supported by our previous observation of prominent TAB2 expression within the EC ^18^ and could explain the high prevalence of valve-defects in patients with *TAB2* defects ^11^.

In both *tak1* and *tab2* mutant zebrafish larvae, we observed significantly reduced trabeculation. Development of the cardiac trabeculae is regulated by a delicate balance between synthesis and degradation of the ECM ^49^. In this regard, the pronounced dysregulation of ECM genes in *tak1* mutants appear to provide the molecular explanation for the reduced trabeculation we observed in *tak1* and *tab2* mutants. Considering the many similarities between our zebrafish mutants and the clinical phenotype in patients, the trabeculation defects observed in *tak1* mutants might warrant increased attention on trabeculation defects among patients with severe *TAK1* variants.

Finally, extracardiac defects are observed among FMD, CSCFS, and THS patients ^11,13,14,19^. Since *tak1* and *tab2* mutants were lethal prior to adult stage, it was not possible to investigate extracardiac defects in detail, but we noted some resemblances with the human syndromes, in the form of craniofacial and pectoral fin anomalies. Thus, the *tak1* and *tab2* mutants appear valid as animal models for FMD, CSCFS, and THS, offering the potential for further advancing our understanding of disease pathomechanisms and as screening models for development of small molecule therapeutics.

In conclusion, our results identify ciliary coordination of TAK1 signaling as an essential mechanism in cardiac development and provide novel insight into the pathomechanisms of human syndromes related to TAK1, TAB2 and PKA-Cα.

## Materials and Methods

### Burden of *TAK1* and *TAB2* variants in CHD patients

Variant burden was calculated using ES data from a case-control dataset of 4,747 CHD cases and 52,881 controls ^20^. Briefly, case and control data were processed and harmonized using the same alignment (BWA v0.7), variant calling (GATK v4.1), variant annotation (VEP API 96) and quality control (Hail v0.2) pipelines. Rare variants (MAF < 0.001) were defined based on its population and cohort-specific allelic frequency. Protein-altering variants (PAVs) include loss-of-function variants and cover the following VEP classifications; ‘missense_variant’, ‘stop_gained’, ‘frameshift_variant’, ‘inframe_deletion’ and ‘splice_region_variant’. The number of individuals with rare (MAF < 0.001, gnomAD v2.1.1 and 3.0.0) synonymous and protein altering variants in *TAK1* and *TAB2* was identified for cases and controls. The variant burden was calculated as the fraction of individuals with variants, divided with the total number of individuals in each group. The statistical significance of differences in variant burden was calculated using Fisher’s exact test.

### Zebrafish maintenance

Embryos were maintained and staged as previously described ^50^ and all experiments were approved and conducted according to licenses and guidelines from the Danish Animal Experiments Inspectorate.

### Zebrafish CRISPR mutants and morphants

The *tak1* (also called *map3k7*, ENSDARG00000020469) and *tab2* (ENSDARG00000021509) CRISPR mutants were generated by injecting gRNA and Cas9 mRNA into 1-cell stage AB wildtype strain zebrafish, acquired from European Zebrafish Resource Center (EZRC). The F0 embryos were raised to adulthood and bred to AB zebrafish line. F1 generation was screened for mutation by colony PCR and Sanger sequencing and founders were bred to *Tg(myl7:GFP, kdrl:mCherry)* line. All mutant analyses were performed on F3-5 embryos, generated by breeding of heterozygous adult fish. As a control either a mix of *+/+* and +/- siblings from +/- adult breeding or +/+ embryos from +/+ (siblings of +/- adults) breeding were used. Genotyping of the larvae was done by PCR (primers are listed in Table 2) and Sanger sequencing. Zebrafish *tak1* and *tab2* morphants were generated by injecting 1 ng of *tak1* (5’ GGAAAGTATTCAAAACTTGCCTTCG 3’) and 0.5 ng of *tab2* (5’ ATCACTCTTGTTCTGAGGAAAGAAG 3’) splice blocking morpholinos respectively to the AB zebrafish line. Standard morpholino (5’ CCTCTTACCTCAGTTACAATTTATA 3’) was used as a control. RT-PCR was performed for testing morpholino efficiency. Primers are listed in Table 2.

### Zebrafish immunostaining, cell counts and assessment of cardiomyocyte proliferation index

For wholemount immunostaining, zebrafish larvae were incubated in 0.2% tricaine for 1 min, fixed in 4% paraformaldehyde (PFA) overnight at 4°C, washed with PBS-T (1X phosphate-buffered saline (PBS) with 0.1% Tween-20), permeabilized with 0.5% Triton X-100 overnight at 4°C and blocked in 5% BSA and 5% goat serum for 2h at RT. Primary and secondary antibodies were added overnight at 4°C, and washed with PBS-T. DAPI staining solution (1 ug/ml) was added for 30 min and larvae were washed with PBS-T prior to imaging. For extracted heart immunostaining, protocol was used as previously described ^51^. Hearts were extracted either manually or using a filter method as described in Lombardo et al ^52^. Zeiss LSM 780 or 980 confocal microscopes were used for imaging. Cell size was measured using Fiji ^53^. All cell counts were performed using the Cell Counter in Fiji. Cardiomyocyte proliferation index was calculated after counting cells in the extracted hearts immunostained with Mef2a/c and PCNA, using the formula: (Mef2^+^ PCNA^+^ cells) / (Mef2^+^ cells). For full list of primary and secondary antibodies used for immunostaining, please see Suppl. Tables 2 and 3, respectively.

### Zebrafish cardiac chamber size, cartilage, fin and eye distance measurements

For cardiac chamber size measurements zebrafish larvae hearts were stopped in 0.2% tricaine and larvae were imaged on 3% methyl cellulose by Zeiss AxioZoom V16. Cardiac chamber area was measured using Fiji ^53^. Alcian blue staining for facial and fin cartilage was performed according to previous protocols with minor alterations ^54^. In brief, embryos were euthanized at 5 dpf before being fixed overnight at 4 °C in 4% PFA. Embryos were then washed with PBS-T and bleached with 3% H_2_O_2_ and 0.5% KOH for 30 minutes. Embryos were carefully washed in PBS-T three times before they were stained with 0.01% Alcian blue/60 mM MgCl_2_/70% ethanol (EtOH) overnight at 4°C. Embryos were then incubated for 10 minutes at RT in 80% EtOH/10 mM MgCl_2_, 50% EtOH/10 mM MgCl_2_, and 25% EtOH/10 mM MgCl_2_ respectively. Further bleaching of the embryos was carried out with 3% H_2_O_2_ and 0.5% KOH for 15 minutes. Washes with 25% glycerol and 0.1% KOH was performed at least twice or until any remaining bubbles from the bleach disappeared. Samples were then stored at 4°C in the dark in 50% glycerol and 0.1% KOH. For cartilage, fin and eye distance measurements, embryos were imaged ventrally on 3% methyl cellulose with a Zeiss AxioZoom V16 microscope. Distances and angles were measured using Fiji as previously described ^53^ and plotted.

### Zebrafish cardiac rate and contractility measurements

Zebrafish larvae heart rates were counted manually under Zeiss Stemi 508 microscope for 15 sec. and then multiplied to get beats/min. For cardiac contractility measurements, Zebrafish larvae were anesthetized in 0.005% tricaine for 3 min and mounted in 0.8% low melting point agarose (Invitrogen). Recordings were performed by Photometrics Prime BSI camera attached to an Olympus BX63 microscope at 100 fps. The % of EF (ejection fraction) and FS (fractional shortening) calculations were done upon ventricular diameter measurements by Fiji as previously described ^55^.

### Zebrafish transcriptome analysis

Sibling *tak1^+/-^* and *tak1^+/+^* adult zebrafish were used for breeding. The phenotypic *tak1^-/-^* larvae were selected from the *tak1^+/-^* breeding; while the *tak1^+/+^*larvae were obtained from the *tak1^+/+^* breeding. At 3 dpf, zebrafish larvae hearts were extracted as previously described ^52^. Three groups of hearts (15 hearts each) were collected from *tak1^-/-^* and *tak1^+/+^* GFP positive larvae. Total mRNA was isolated using Qiagen Rneasy micro kit according to manufacturer’s protocol. Bulk RNAseq was performed by BGI, China. Remaining tissue was collected from the larvae for verifying the genotype by Sanger sequencing. The sequencing chromatogram from phenotypic larvae revealed clear peaks with a single bp deletion as expected from *tak1^-/-^* zebrafish. Differentially expressed genes were identified as genes with statistically significant difference in expression between *tak1^-/-^* and *tak1^+/+^* samples (q-value <0.05). Human orthologues of zebrafish genes were selected using HGNC Comparison of Orthology Predictions (HCOP) (https://www.genenames.org/tools/hcop/). GO annotation analysis was performed using WebGestalt (https://www.webgestalt.org/) ^56^. Gene enrichment was determined by calculating overlap between gene lists. Significant overlap was calculated using hypergeometric statistics (http://nemates.org/MA/progs/overlap_stats.html). A representation factor was calculated as the number of overlapping genes, divided by the expected number of overlapping genes drawn from two independent groups: RF=x/((n*D)/N), where x=number of overlapping genes, n=genes in group 1, D=genes in group 2, N=genes in genome (20,000).

### RPE-1 cell culture, stimulation assay and transfection

RPE-1 (human telomerase reverse transcriptase-immortalized retinal pigmented epithelial-1) cells were cultured in T75 flasks at 37°C and 5% CO2 in DMEM (Gibco) supplemented with 10% fetal bovine serum (FBS, Gibco) and 1% penicillin-streptomycin (Sigma-Aldrich) and passaged with 1% Trypsin (Sigma-Aldrich) when the cells reached 80% confluency. RPE-1 cells were seeded as 2,000 cells/cm^2^ in DMEM (+/+) media in culture dishes. After 24 hours, cells were washed with 1x PBS and serum starved in serum depleted DMEM for 48 hours. The cells were then stimulated with TGFB1 (R&D Systems, #240-B) or BMP2 (R&D Systems, #355-BM) ligand at a final concentration of 2ng/mL and 100ng/mL respectively as previously described ^57^. The cells were then fixated with 4% PFA for IFM or lysed with SDS for western blot. The RPE-1 cells were transfected with plasmid constructs p3XFLAG-TAK1-WT, p3XFLAG-TAK1-P485L, p3XFLAG-TAK1-G168R which comprised wildtype TAK1 and two FMD patient mutations, respectively. Transfection was achieved using Lipofectamine300® Transfection Kit (Invitrogen, L3000-015, 2201452). Briefly, RPE-1 cells were seeded as 100,000 cells/well in 6-well plates with DMEM medium supplemented with 10% FBS and 1% penicillin-streptomycin for 24 hours. The cells were then serum starved for 48 hours followed by incubating with transfection mixture and 1.5 μg of respective plasmid DNA for 8 hours. Cells were then analyzed for IFM or western blot.

### P19CL6 mouse stem cell culture, cardiomyogenesis and TAK1 inhibition

CRISPR-Cas9 gene editing was used to generate P19CL6 cell mutants. For *Tak1*, an 85 bp deletion was introduced in *Tak1* that induced exon 2 skipping resulting in *Tak1* homozygote KO clone. For *Tab2*, a heterozygote *Tab2* KO clone was generated by inducing a 58 bp deletion on allele A leading to a premature stop codon at nucleotide 418. Additionally, a 10 and 25 bp deletion was introduced on allele B causing a premature stop codon at nucleotide 298. The P19CL6 wildtype and *Tak1* and *Tab2* mutants were cultured in T25 flask at 37°C and 5% CO2 in MEM alpha medium (Gibco) supplemented with 10% FBS (Gibco) and 1% Penicillin-Streptomycin (Sigma-Aldrich) and passaged with 1% Trypsin (Sigma-Aldrich) when reached at 80% confluency. For cardiomyocyte differentiation, the P19CL6 cells were seeded with a density of 1000cells/cm^2^ with an induction by 1% DMSO in the culture medium. Differentiation assay was performed as previously described ^33^. For TAK1 inhibition assay, P19CL6 cells were seeded as 4800 cells/mL in the presence of either 10 uM Takinib (ToCris #6430) or 500 nM (5Z)-7-oxozeanol (5ZO) (ToCris #3604) or the vehicle and set to differentiate in the presence of 1% DMSO.

### Immunofluorescence microscopy analysis of RPE-1 and P19.CL6 cells

RPE-1 and P19CL6 cells were grown on coverslips and washed with 1x PBS, fixed in 4% formaldehyde for 15 and 25 minutes, respectively, permeabilized in 0.2% Triton X-100 at room temperature for 12 minutes and blocked with 2% BSA for 30 minutes. Primary antibodies (Suppl. Table 2) were incubated at 4°C, overnight and secondary antibodies (Suppl. Table 3) at room temperature for 45 minutes followed by 30 seconds incubation with DAPI. Images were visualized using an Olympus BX63 upright microscope with a DP72 digital camera. Olympus CellSens dimension software was used to measure the fluorescent intensities at the cilium-centrosome axis. The mean fluorescence values at the cilium-centrosome axis were set relative to the fluorescence values in background areas of the cytosol. Images were processed for publication using Adobe Photoshop CS6 (version 13.0) and data were presented using GrapPad Prism software (version 9). GATA4 and α-actinin quantification was performed by findings regions filled with nuclei via the DAPI channel, changing to the channel of interest and then quantifying the amount of positive nuclei or the total fluorescent intensity of the frame respectively.

### Seeding, differentiation and immunofluorescence microscopy analysis of gastruloids

mESCs required for gastruloid formation were procured from an 80% confluent T25 gelatin-coated flask. The old medium was aspirated followed by a wash in 10 mL Gibco™ pH7.4 PBS. Hereafter, the cells were trypsinized with Gibco™ TrypLE Express (#12605010) and incubated for 5 minutes at 37°C, 5% CO2. Trypsin was neutralized with the addition of 7 mL Gibco™ DMEM medium (#11965092) + Trypsin Neutralization solution (#R002100). The cells were then centrifuged at 170 g at room temperature for 5 minutes. The pellet was washed twice by adding 10 mL PBS, then centrifuging the cells for 5 minutes at 170 g. The cells were resuspended in 5 mL N2B27 medium (Appendix 1.4). Using a kova-slide (Pierron, 13367.20), the cells were counted and seeded to 7500 cells/mL, whereafter 40 μL was added to each well of a 96 well plate, making the final cell-count per well approximately 300. The cells were then incubated at 37°C, 5% CO2 for two days. Following gastroloid seeding, 150 μL of fresh N2B27 medium + 3 μM CHIR99021 was added to the cells that were then incubated at 37°C, 5% CO_2_ for one day. On day 3, 120 μL of the old medium was removed and replaced with further 150 μL of fresh N2B27 medium. At the beginning of day 4, gastruloid elongation was observed. 150 μL of the old medium was replaced with 150 μL of fresh N2B27+++ medium whereafter the 96-well plates were placed on an orbital shaker at continuous shaking at 100 rpm at 37 C°, 5% CO_2_. Fixed gastruloids were serially dehydrated in EtOH/PBS and embedded in paraffin. Lateral sections were collected for analysis. Sections were de-paraffinized with several washes in xylene, followed by serial gradient rehydration in EtOH. Sections were boiled in citrate buffer (pH 6.0) or Tris-EGTA (TEG) buffer (pH 8.0) for approximately 20 minutes and cooled. Blocking buffer (DAKO Real Antibody Diluent, #S2022) was added to the samples for 30 minutes before the sections were incubated with primary antibodies (Suppl. Table 2) diluted in blocking buffer overnight at 4°C. Sections were then washed with 1X TBS (Tris-buffered saline) before incubating for 45 minutes in secondary antibodies (Suppl. Table 3) and DAPI diluted in blocking buffer. Washes with 1X TBS was performed again before a final wash in dH_2_O. Images were taken on an Olympus BX63 upright microscope as described for RPE-1 and P19.CL6 cells.

### SDS-PAGE and western blotting analysis

RPE-1 and P19CL6: cells were washed in 1x PBS and lysed with 1% SDS (1% SDS, 1M Tris-HCL (pH 7.5)) and EBC (4 M NaCl, 1 M Tris HCL (pH 7.5), 0.5 M EDTA, 0.5% NP-40) lysis buffers, respectively. Lysates was homogenized by sonication and centrifuged to collect the supernatant. Zebrafish larvae: approximately 15, 3 dpf larvae were dechorionated manually and yolk was removed in deyolking buffer (55mM NaCl, 1.8mM KCl, 1.25 mM NaHCO3) by pipetting and shaking using a thermomixer. Larvae were pelleted at 3,000 rpm for 1 min at 4°C and washed twice with wash buffer (110 mM NaCl, 3.5 mM KCl, 2.7mM CaCl2, 10 mM Tris (pH 8.5)). Following the last wash, supernatant was removed and 80 μl of 1% SDS lysis buffer supplemented with protease inhibitor was added. Pellet pestle motor (VWR, # 47747-370) was used to dissociate larvae. Lysates were stored in −80°C until use. For all lysates, protein concentrations were determined using BioRad DC Protein Assay, OD measured at 750 nm using a Beckmann Coulter DU spectrophotometer. SDS-PAGE and western blot was performed as previously described ^58^. Blots were developed in FUSION-FX chemiluminescence system. Band intensities were analyzed by densitometric scanning using UN-SCAN-IT 6.1 software (Silk Scientific). Please see Suppl. Tables 2 and 3 for primary and secondary antibodies, respectively.

### Statistical analyses

Unless otherwise mentioned, statistical analysis was carried out using the student’s t test for comparing the variation between two groups. All statistical calculations were performed on n=3 or more. Significance levels were as follows: ^∗^p < 0.05, ^∗∗^p < 0.01, and ^∗∗∗^p < 0.001, ns: not significant.

## Acknowledgements

We are grateful to the patients and their families for their participation in the project. We would like to thank Lillian Rasmussen and Søren Lek Johansen for excellent technical assistance and we are grateful to Simon Holst Bekker-Jensen for reagents. We would further like to thank Paula Lillo López, Signe Grue Andreassen, Emilie Damm Garly, Julie Linnea Hammer Dragheim, Osama Jamal Bin Amir Hussain, Magnus Aalborg Nielsen, Jeppe Theisen Pedersen and Malou Maria Nielsen for their assistance in experimental procedures and data collection. The study was supported by grants from the Danish Heart Foundation, Børnehjertefonden, Novo Nordisk Foundation, Carlsberg Foundation, European Commission Horizon 2020 research and innovation programme Marie Skłodowska-Curie Innovative Training Networks (grant agreement No. 861329), Independent Research Fund Denmark (grant no. 3103-00177B), the Novo Nordisk and Novozymes Scholarship Program, Læge Sophus Carl Emil Friis og hustru Olga Doris Friis’ Legat, Direktør Jacob Madsens og Hustru Olga Madsens Fond, Fonden til Lægevidenskabens Fremme, Helge Peetz og Verner Peetz og hustru Vilma Peetz Legat and Islamic Development Bank (IsDB).

## Author Contributions

LAL and STC conceived the study. LAL, STC, LBP, FL and M-PH supervised experiments and analyses. CD, DB, OKT, YA, LAL and STC wrote the manuscript with input from all authors (EA, LJ, PMT, JBM, MSH, KA, LB, MD, JLF, EMW, SPR, LBP, LA, FL, M-PH). EA, MP-H and LAL performed human genetic analyses. CD, DB, OKT, YA, PMT, KA, LB, MD and JLF performed zebrafish experiments and analyses. CD and LAL performed RNAseq experiments and analyses. YA, OKT, LJ, DB, JLF, MSH, JBM and LA performed experiments and analyses on cell cultures and gastruloids.

## Declaration of interests

The authors declare no competing interests.

---

**Suppl. Figure 1.**
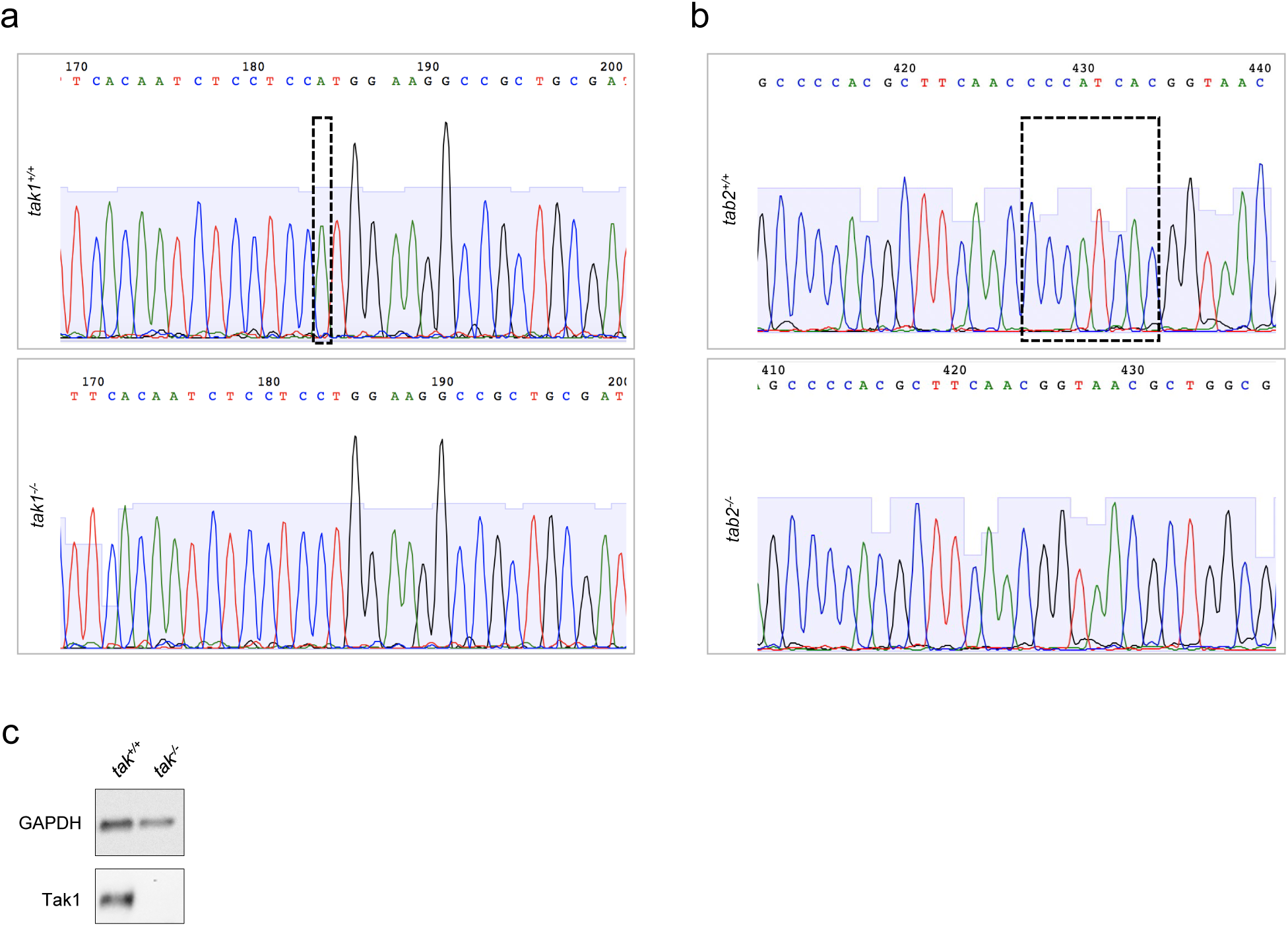
CRISPR-Cas9 editing in zebrafish *tak1* and *tab2*. **a, b.** Sanger sequencing chromatograms displaying *tak1^-/-^* (a) and *tab2^-/-^* (-b) mutant zebrafish sequences in comparison to *tak1^+/+^*and *tab2^+/+^* wildtype sequences. Dashed boxes mark deleted nucleotides in the mutant zebrafish. **c.** validation of Tak1 knockout in *tak1^-/-^*mutant zebrafish by western blotting analysis.

**Suppl. Figure 2.**
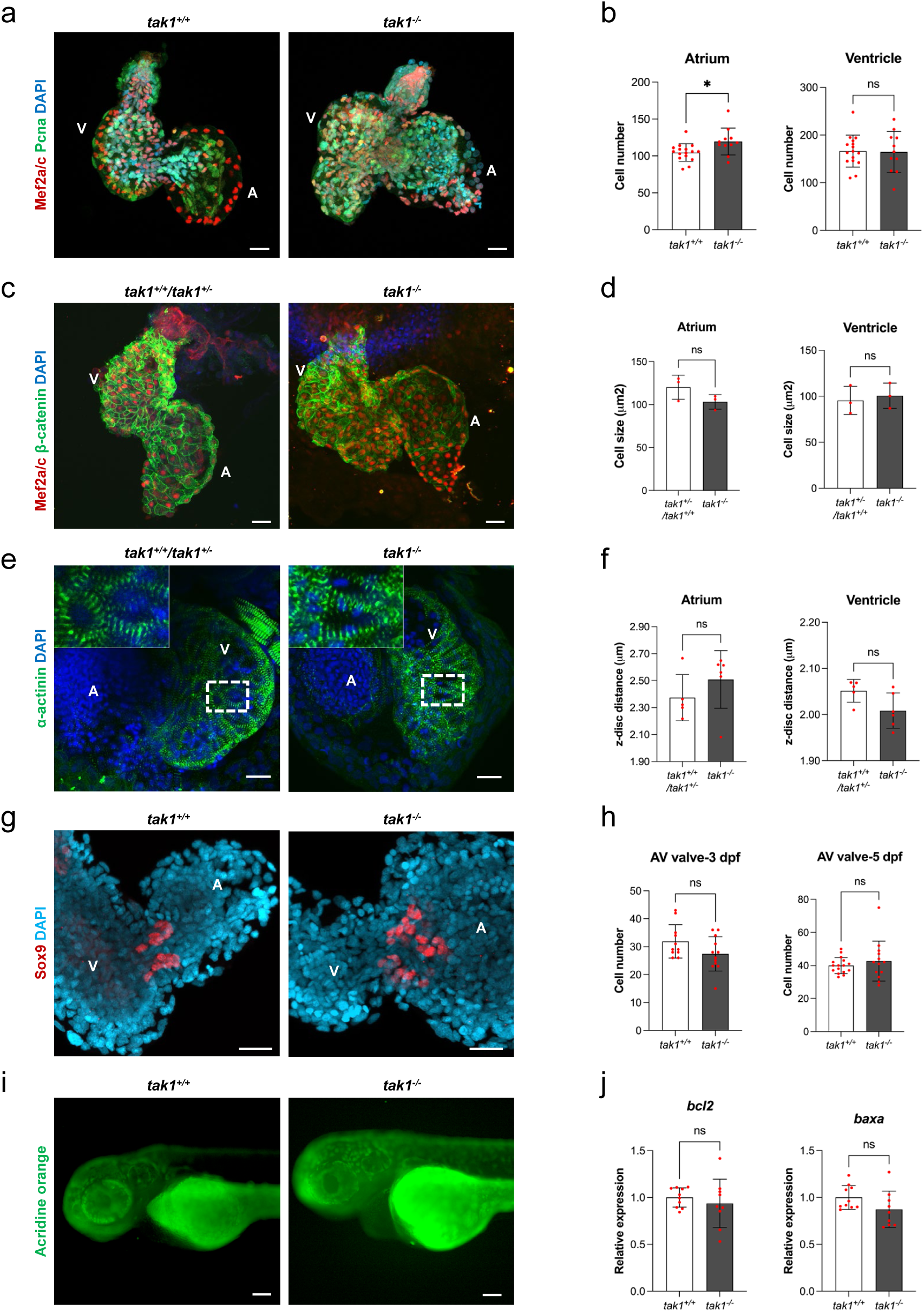
Cardiac phenotype analysis of *tak1* mutant. **a**. IFM with Mef2a/c (red) and Pcna (green) antibodies in 3 dpf *tak1^+/+^* and *tak1^-/-^* extracted zebrafish hearts. Scale bars, 20 μm. **b**. Quantification of number of cardiomyocytes (Mef2a/c-positive cells) in atrium and ventricle of 3 dpf *tak1^+/+^*and *tak1^-/-^* zebrafish hearts. **c**. IFM with β-catenin (green) and Mef2a/c (red) antibodies in 3 dpf *tak1^+/+^*/*tak1^+/-^* and *tak1^-/-^* extracted zebrafish hearts. Scale bars, 20 μm. **d**. Quantification of cell size in atrium and ventricle of 3 dpf *tak1^+/+^*/*tak1^+/-^* and *tak1^-/-^* zebrafish hearts. Each dot represents an average of 10 cells in a larvae. **e**. Wholemount IFM with α-actinin (green) antibody marking z-discs in the 3 dpf *tak1^+/+^*/*tak1^+/-^* and *tak1^-/-^* zebrafish hearts. Scale bars, 20 μmm. **f**. Quantification of z-disc distance in the atrium and the ventricle of 3 dpf *tak1^+/+^*/*tak1^+/-^*and *tak1^-/-^* zebrafish hearts. Each dot represents an average of 10 sarcomere units in a larvae. **g**. IFM with Sox9 antibody (red) in 3 dpf *tak1^+/+-^*and *tak1^-^* extracted zebrafish hearts. Scale bars, 20 μm. **h**. Quantification of number of mesenchymal cells (Sox9 positive cells) in AV valves of *tak1^+/+-^* and *tak1^-^* zebrafish hearts at 3 dpf and 5 dpf. **i**. Acridine orange staining (green) of 3 dpf *tak1^+/+-^*and *tak1^-^* zebrafish showing apoptotic cells. Scale bars, 0.1 mm. **j**. Relative expression of bcl2 (left) and baxa (right) in 3 dpf *tak1^+/+-^* and *tak1^-^* extracted zebrafish hearts. *: P<0.05, ns: not significant. Nuclei were stained with DAPI (blue).

**Suppl. Figure 3.**
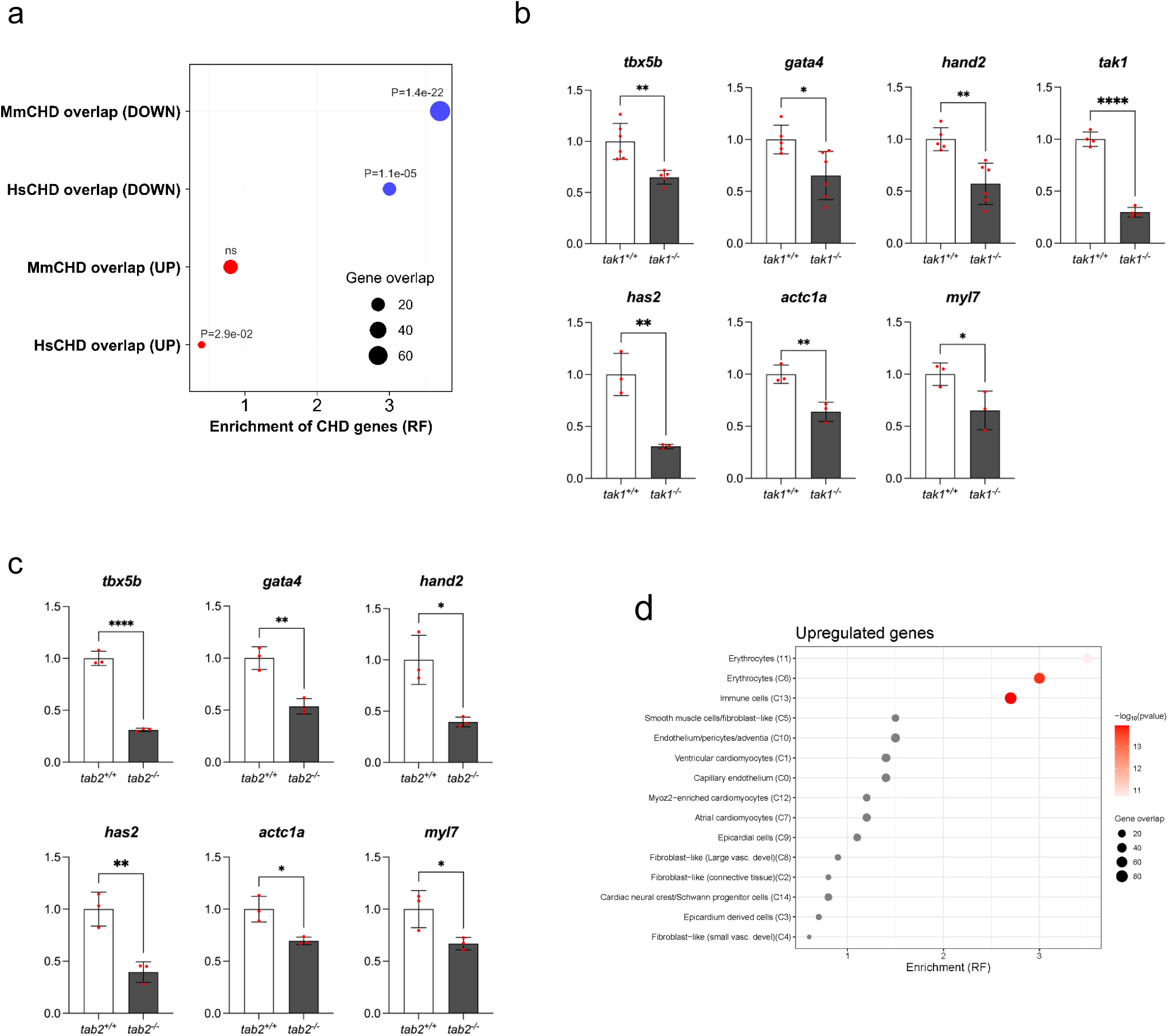
Phenotype of *tab2* mutant. **a.** Gross morphology of 5 dpf *tab2^+/+^* and *tab2^-/-^* zebrafish larvae highlighting pericardial edema (black arrow) and protruding mouth (open arrow). Scale bars, 0.2 mm. **b.** Survival of *tab2^+/+^*, *tab2^+/-^* and *tab2^-/-^* zebrafish larvae. **c.** Upper panels: wholemount IFM of transgenic (*Tg[myl7:GFP]*) *tab2^+/+^* and *tab2^-/-^* zebrafish hearts with GFP (green) and S46 (red) antibodies at 5 dpf. A: atrium, V: ventricle. Lower panels: wholemount IFM of transgenic (*Tg[myl7:GFP]*) *tab2^+/+^* and *tab2^-/-^*zebrafish hearts with GFP antibody (green) at 6 dpf. Nuclei were stained with DAPI (blue). Ventricular trabeculae are indicated with arrows and trabeculation defects with asterisks. Scale bars, 20 μm. **d.** Quantification of normalized ventricle area, atrium area, heart rate, fractional shortening (%) and ejection fraction (%) at 5 dpf. **e.** IFM with Sox9 antibody (red) in 5 dpf *tab2^+/+^* and *tab2^-/-^* extracted zebrafish hearts. Nuclei were stained with DAPI (blue). Scale bars, 20 μm. **f.** Quantification of number of mesenchymal cells (Sox9-positive cells) in AV valves of *tab2^+/+^* and *tab2^-/-^*zebrafish hearts at 5 dpf. **g.** Bright-field images of *tab2^+/+^/tab2^+/-^*(upper panels) and *tab2^-/-^* (lower panels) larvae for (i) Me’kel’s-palatoquadrate (M-PQ) angle measurements of cartilage with Alcian blue staining at 5 dpf. Black lines show the measured angle in larvae. (ii) Measurement of fin length with Alcian blue staining at 5 dpf. Black line shows measured fin length. (iii) Eye distance measurement at 7 dpf. Black line shows the measured eye distance in larvae. Scale bars, 0.1 mm. **h.** Quantification of measurements obtained from i-iii. **: P<0.01, ***: P<0.001, ****: P<0.0001, ns: not significant.

**Suppl. Figure 4.**
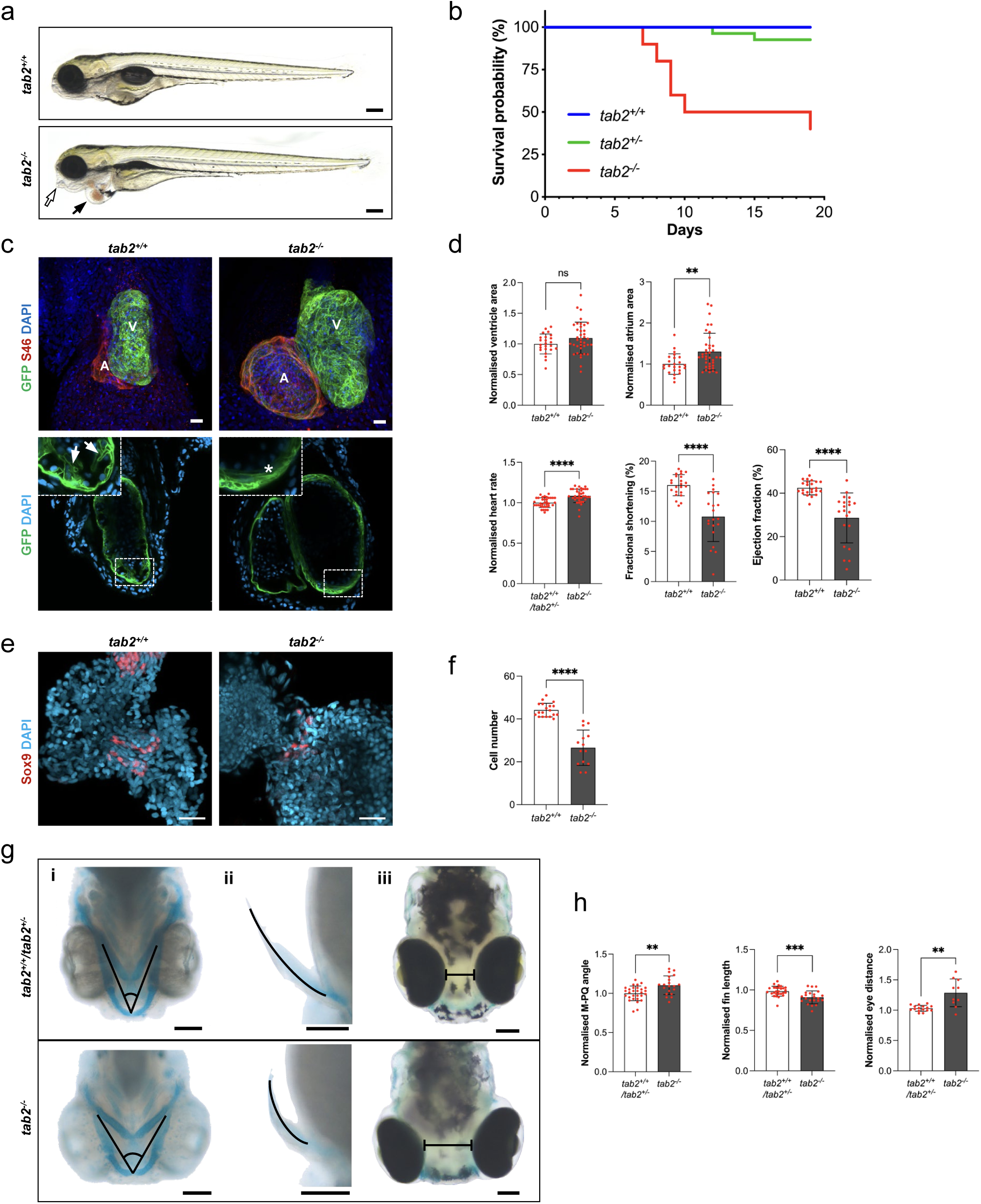
Morpholino knockdown of zebrafish *tak1* and *tab2*. **a.** Illustration of the *tak1* and *tab2* splice blocking morpholino target sites and PCRs showing their efficiency. **b.** Phenotype assessment of *tak1*-MO and *tab2*-MO injected embryos at 2 dpf and the rescue assays by corresponding mRNA co-injections. **c.** Quantification of the cardiac phenotypes in *tak1*-MO and *tab2*-MO injected embryos at 2 dpf at optimal doses. **d.** Quantification of the cardiac phenotypes upon co-injection of *tak1*-MO and *tab2*-MO at sub-efficient doses. Statistical analysis was performed via two-way ANOVA. *:P<0.05, **: P<0.01, ***: P<0.001, ****: P<0.0001

**Suppl. Figure 5.**
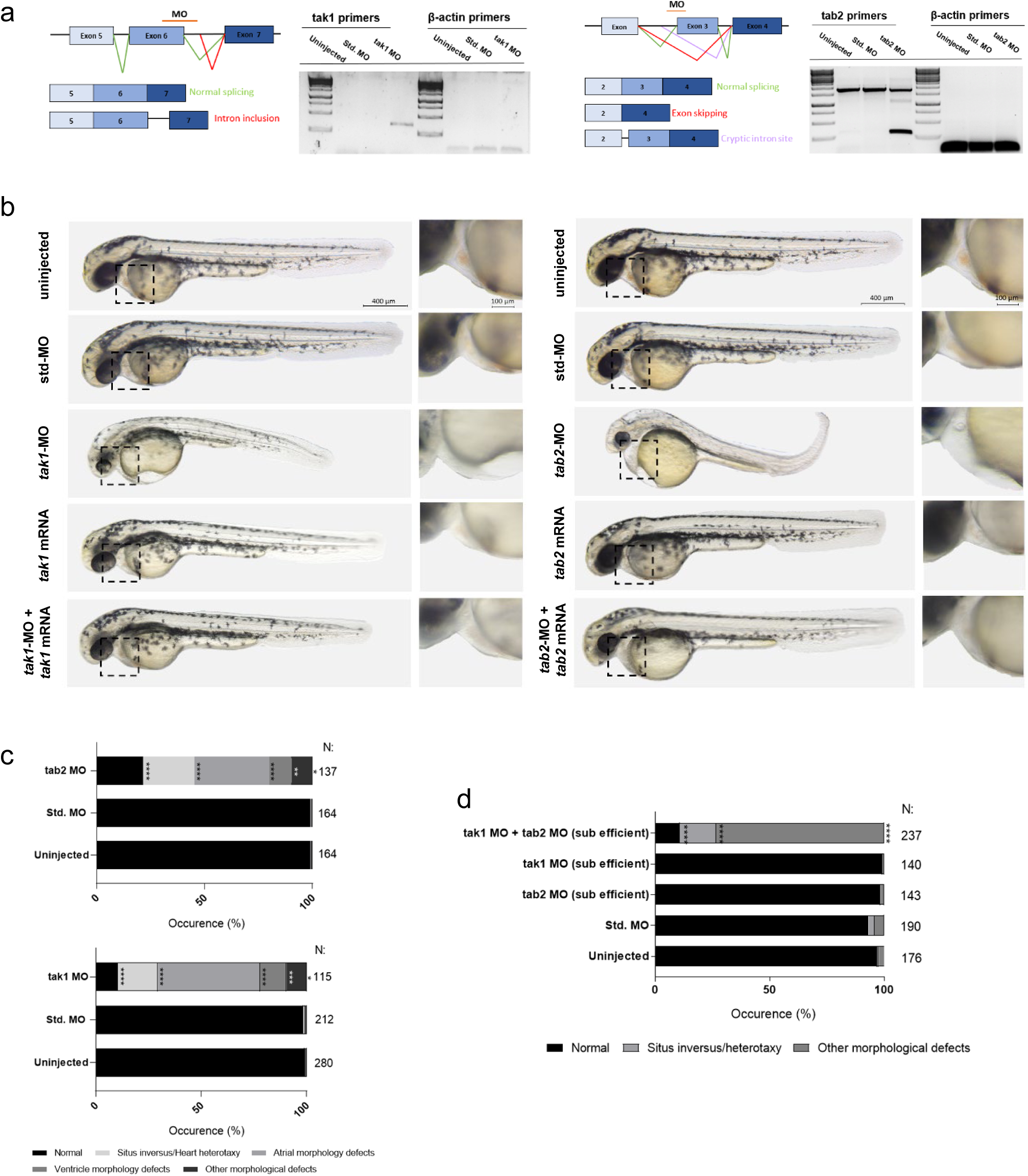
RNA sequencing and RT-qPCR. **a.** Enrichment of known CHD genes within 474 down- and 712 upregulated genes (blue and red circles, respectively). Enrichment is shown as representation factor (RF). A hypergeometric distribution was used to test the significance. MmCHD: a list of 832 genes causing heart defects in mouse models ^22^. HsCHD: a list of 377 genes causing CHD in human patients ^21^. **b, c.** RT-qPCR of differentially expressed genes found in RNAseq between (b) *tak1^+/+^* and *tak1^-/-^* extracted hearts (RT-qPCR for *tak1* was performed on the whole larvae) at 3 dpf and (c) *tab2^+/+^*and *tab2^-/-^* extracted hearts at 5 dpf. The samples used for RT-qPCR verification were independent from the samples used for RNAseq. *:P<0.05, **: P<0.01, ****: P<0.0001. **d.** Enrichment of upregulated genes in the expression signature of fifteen cell-types in 6.5-7 PCW human developing hearts ^23^. Enrichment is shown as representation factor (RF). A hypergeometric distribution was used to test the significance. Non-significant enrichment is marked with grey circles.

**Suppl. Figure 6:**
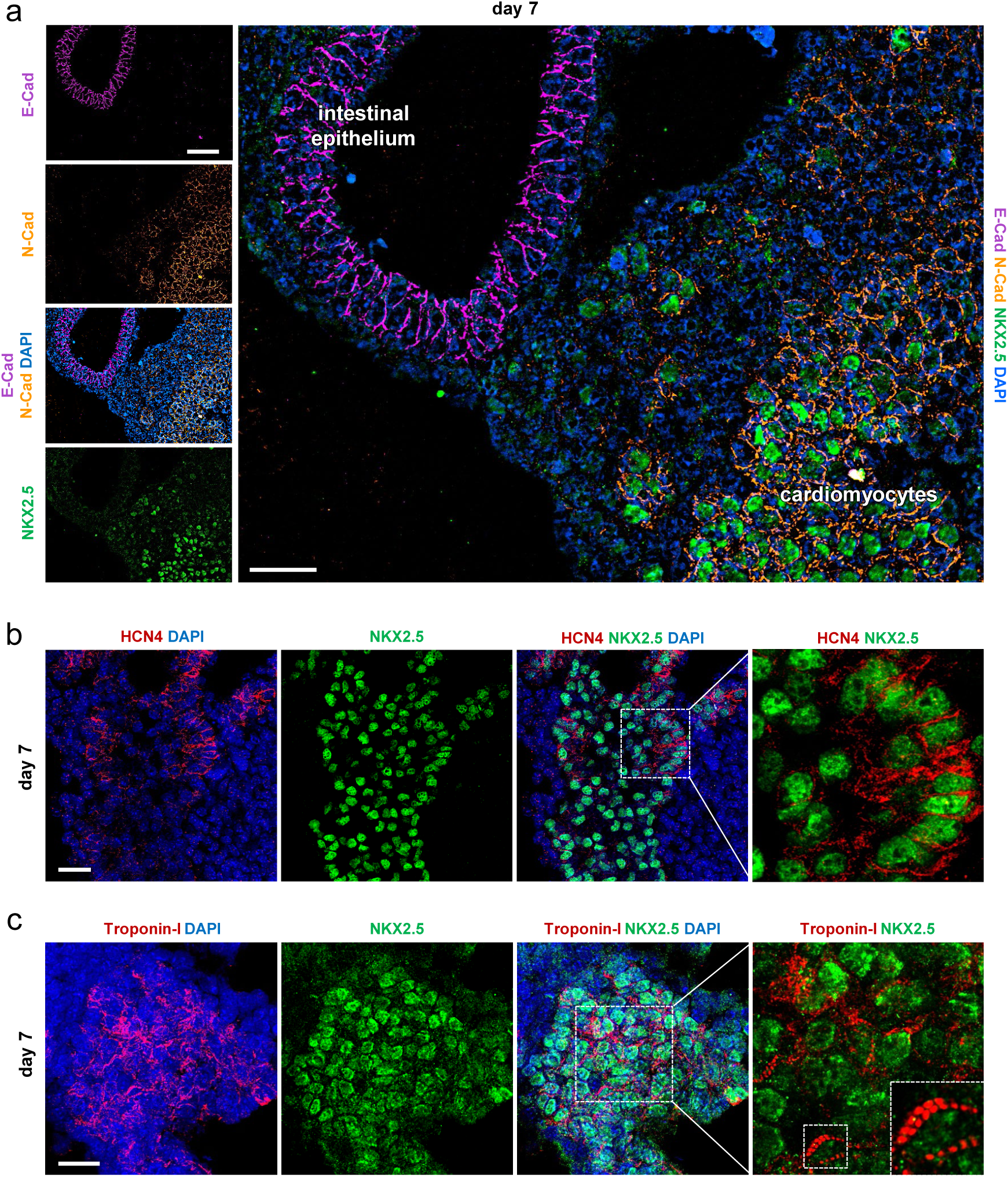
Characterization of day 7 gastruloids. **a.** Representative IFM image indicating the expression of cardiac junction protein N-Cadherin (pseudocolored orange) in NKX2.5 positive cells/cardiomyocytes (green) and the epithelial cadherin, E-Cadherin (pseudocolored magenta) in intestinal epithelium of day 7 gastruloids. Small image scale bar, 40 µm, large image scale bar, 20 µm. **b, c.** Representative IFM images of day 7 gastruloids showing HCN4 (red, marks FHF) (b) and Troponin-I (red) (c) in NKX2.5 (green) positive cells. Scale bars, 20 µm.

**Suppl. Figure 7:**
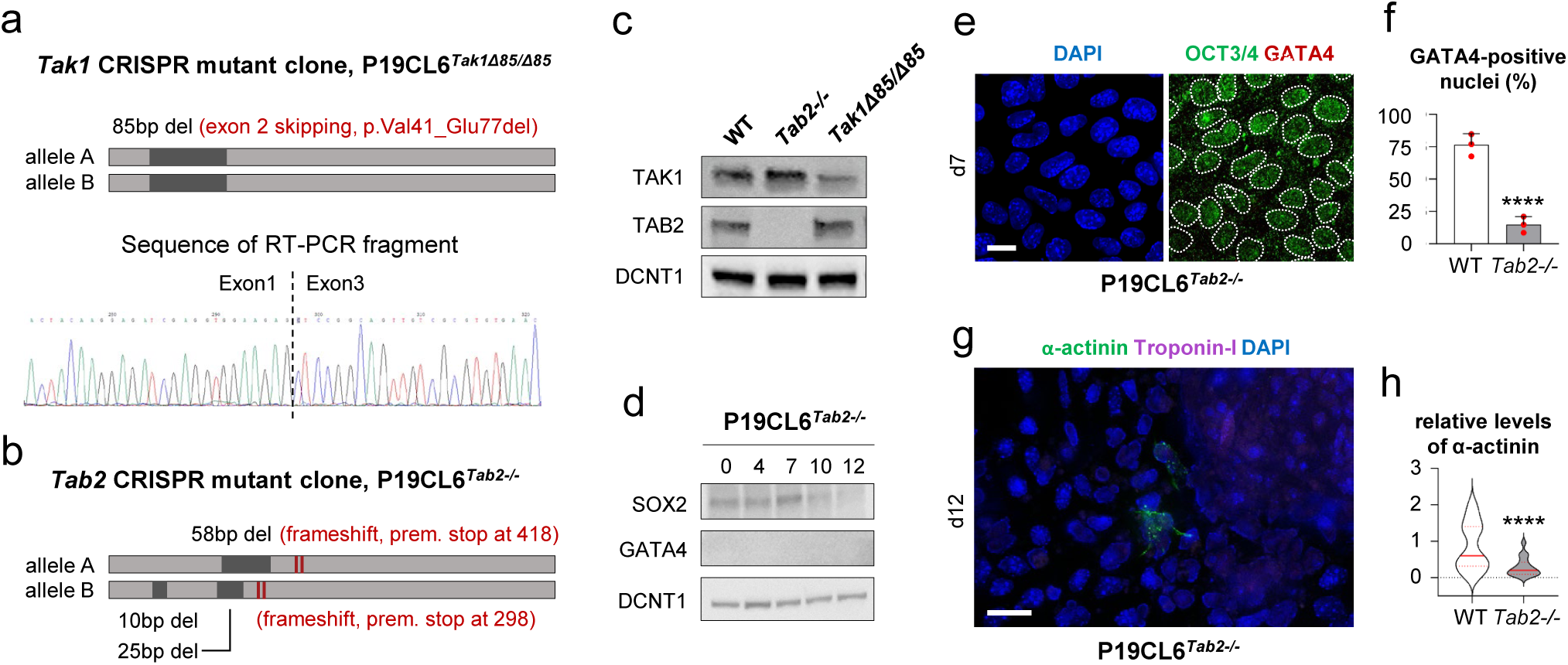
CRISPR-Cas9 editing in P19CL6 cells and differentiation phenotype in *Tab^-/-^* cells. **a, b.** Graphical illustration of *Tak1* CRISPR mutant clone P19CL6*^Tak1Δ^*^85*/Δ*85^ (a) and *Tab2* CRISPR mutant clone P19CL6*^Tab^*^2*-/-*^ (b) showing deletions (black) and predicted premature stop codons (red lines). The deletion in P19.CL6*^Tak1Δ^*^85*/Δ*85^ cause aberrant splicing (skipping of exon 2), which results in a deletion (p.Val41-Glu77del). **c.** WB analysis of P19CL6^WT^, P19CL6*^Tab^*^2^*^-/-^* and P19CL6*^Tak1Δ^*^85*/Δ*85^ cells using antibodies against TAK1, TAB2 and DCTN1. **d.** WB analysis of DMSO treated P19.CL6*^Tab^*^2^*^-/-^* cells using antibodies against SOX2, GATA4 and DCTN1. **e.** Representative IFM images of expression of OCT3/4 (green) and GATA4 (red) at day 7 of DMSO stimulation in P19CL6*^Tab^*^2^*^-/-^* cells. Scale bar, 10 μm. **f.** Percentages of GATA4-positive nuclei in P19CL6^WT^ and P19CL6*^Tab^*^2*-/-*^ at day 7. ****P<0.0001. **g**. α-actinin (green) and Troponin-I (pseudocolored magenta) at day 12 of DMSO stimulation in P19CL6*^Tab^*^2*-/-*^ cells. Scale bar, 20 μm. **h.** Relative levels of α-actinin in P19.CL6^WT^ and P19.CL6*^Tab^*^2^*^-/-^*cells at day 12. ****P<0.0001.

## Notes

### Competing Interest Statement

The authors have declared no competing interest.

